# Deciphering colchicine like actions of clerodin in terms of microtubule destabilization based mitotic abnormalities, G2/M-phase arrest, and plant polyploidy

**DOI:** 10.1101/2020.12.27.424481

**Authors:** Sujit Roy, Lalit Mohan Kundu, Gobinda Chandra Roy, Manabendu Barman, Tista Chakraborty, Poulomi Ghosh, Ria Das, Debabrata Tripathy, Nanda Singh, Sagnik Banerjee, Atanu Bhattacharjee, Amit Pal, Anupam Chatterjee, Sanjib Ray

## Abstract

Clerodin (C_24_H_34_O_7_), a clerodane diterpenoid, is a bitter principle of *Clerodendrum viscosum*. The present study aimed to decipher colchicine-like actions of clerodin in terms of microtubule destabilization based mitotic abnormalities, G2-M arrest, and plant polyploidy. Purified clerodin showed increased metaphase frequency in Human Peripheral Blood Lymphocytes (HPBLs), Human Embryonic Kidney cells (HEK-293), and *Allium cepa* root apical meristem cells. Both squashed slide of the onion root tip and flow cytometric analysis of radish protoplast revealed a significantly increased frequency of polyploid cells. Flow cytometric analysis showed an increase in frequencies of G2-M in MCF-7 cells from 6.10 to 16.25% after clerodin (200μg/mL) treatment for 24 h. Confocal microscopy imaging of tubulin in clerodin-treated MCF-7 cells revealed microtubule destabilization. Molecular docking and LIGPLOT analysis indicate that clerodin interact in the colchicine binding site, including, single hydrogen bond with Asn 101 of α-tubulin. In summary, our experimental data revealed that clerodin has metaphase arresting, microtubule destabilization, and polyploidy inducing ability similar to colchicine. Molecular docking analysis revealed for the first time that clerodin and colchicine interact at the common site of tubulin residue indicating a common mechanism of action. The results also indicate similar cytotoxic potentialities of both clerodin and colchicine even though they belong to different chemical groups. Thus, clerodin may be used in place of colchicine as a plant polyploidy inducing agent in plant breeding programs in Agriculture.

## 1. Introduction

Plant-derived anticancer agents such as vinblastine, vincristine, camptothecin derivatives, topotecan and irinotecan, etoposide, epipodophyllotoxin derivative, and Taxol (Paclitaxel) are under clinical practice nowadays (Haqueet al. 2016). Different extracts from *Clerodendrum viscosum* Vent. (Family: Lamiaceae) is used as an antidote of snakebite, anthelmintic (Saroj 2016), analgesic and anticonvulsant, antioxidant, antimicrobial, hepatoprotective, wound healing and antidiarrheal (Bhattacharjee et al. 2011), hypoglycemic (Hossain et al. 2014, Chandrashekar et al. 2012), cytotoxic, anthelmintic (Rahman et al. 2013), negative impact on the growth and germination of weeds in an agroecosystem (Devi et al. 2013, Qasem et al. 2001), insect repellent and insecticidal activity (Muh et al. 2014). Recently, we have shown a colchicine-like antiproliferative and metaphase arresting potentials of the aqueous extract of *C.viscosum* leaf (Ray et al. 2013, Kundu et al. 2016). All these results indicate that the clerodane diterpenoids are the major bioactive components of *C. viscosum* (Ray et al. 2013, Kumar et al. 2018, Abbaszadeh et al. 2012). Increased attention has been drawn to clerodane diterpenoids like clerodin because of insect antifeedant role against economically important insect phytophagous pests. Ajugarins I and IV are well known for their insect antifeedant properties (Kubo et al. 1979, Kubo et al. 1982). Ajugarin IV also displays insect growth regulating activity, as do 3-epi-caryoptin (Pereira et al. 1990) and the 19-nor-clerodanes cis-and trans-dehydrocrotonin. Fungicidal activity against plant pathogenic fungi has been reported for clerodin and the related jodrellins A and B (Cole et al 1991).

Clerodin, the first member of the clerodane diterpenoid series and one of the bitter principles of the Indian bhat tree, *Clerodendrum infortunatum*, was first isolated by Banerjee (Barton et al. 1961). Extraction of the ground leaves and twigs of *C.infortunatum* gave a crude solid, from which three crystalline compounds were isolated, of them clerodin was present in major amount. From the X-ray crystallography analysis, the molecular formula of clerodin was established as C_24_H_34_O_7_ (Barton et al. 1961). Growth inhibitory activity against *Helicoverpa armigera* (cotton boll-worm) was exhibited by Clerodin,15-methoxy-14,15-dihydroClerodin, and 15-hydroxy-14,15-dihydroClerodin (Abbaszadeh et al. 2012).

Several clerodane diterpenes also possess various pharmacological activities including anti-ulcer, cytotoxic, anti-inflammatory, antiparasitic, and antibacterial activities (Li et al. 2016). Our previous results indicate, leaf aqueous extract of *C. viscosum* has metaphase arresting, mitotic abnormalities inducing, and antiproliferative activities (Ray et al. 2013, Kundu et al. 2016). Subsequently, bioassay-guided fractionation analysis indicates that clerodane diterpenoids-rich fractions act as spindle poison (Roy et al. 2020).

Colchicine, an alkaloid extracted from *Colchicum autumnale* and *Gloriosa superb*, is used to treat rheumatic arthritis (Ade et al. 2010). It inhibits neutrophil activity and thus acts as an anti-inflammatory agent, relieves the pain from gout outbreaks (Van et al. 2014). Colchicine arrests cells at pro-metaphase by inhibiting spindle formation leading to abnormal separations of chromatids and induction of polyploidy in plants (Salmon et al. 1984, Caperta et al. 2006) and it is now widely used in modern agricultural plant breeding programs to induce plant polyploidy (Ade et al. 2010). The polyploid plants exhibit various advantageous characters like the increased size of organs, drought tolerance, pest resistance, blooming time, ability to survive in the harsh environment, and synthesis of more secondary metabolites than their diploid parents (Manzoor et al. 2019). The wide application of the polyploidy inducing property of colchicine led to increasing not only the ornamental property of plants but also the yield of vegetative crops in agriculture by overcoming hybridization barrier, restoring fertility, developing pest resistance and stress tolerance activity due to an increase in secondary metabolites productions like total phenolics, flavonoids, *etc*. (Manzoor et al. 2019). Here, pure clerodin crystals (95.84%) extracted from *C. viscosum* were tested to analyze their metaphase arresting, microtubule destabilization based mitotic abnormalities, and polyploidy inducing effects using the suitable experimental models from both plant and animal tissues. For colchicine like pro-metaphase arresting activity analysis, Human Peripheral Blood Lymphocytes (HPBLs), Human Embryonic Kidney cells (HEK-293), and *Allium cepa* root apical meristem cells were used. Microtubule destabilization and G2/M arresting activities of clerodin were analyzed through a flow cytometer and confocal microscope on breast cancer cell line MCF-7. Clerodin mediated microtubule destabilization and colchicine induced mitotic abnormalities, and polyploidy were analyzed in onion root tip cells and radish protoplasts. Moreover, a comparative account of molecular docking of colchicine and clerodin with microtubule subunit was performed to explore their binding pockets and interacting residues. The novelty of the present study is that we demonstrate colchicine like actions of clerodin in terms of microtubule destabilization leading to plant polyploidy.

## 2. Materials and methods

### 2.1. Chemicals

Orcein, glacial acetic acid, and methanol were obtained from Merck, Germany. Colchicine, Triton X-100, mannitol, sucrose, pectinase, cellulase, and paraformaldehyde were obtained from Himedia, India. RPMI 1640, DMEM, FBS, Penicillin-streptomycin, and 0.25% Trypsin-EDTA (1X) was obtained from Invitrogen-Gibco. RNase A, Propidium Iodide, and DAPI/Antifade solution were purchased from Sigma-Aldrich, USA. Anti-tubulin antibody and Goat anti-mouse IgG-FITC used were obtained from Santa-Cruz, The USA. Clerodin (95.84% pure needle-shaped crystals, Figure 1) was isolated from leaf aqueous extract of *Clerodendrum viscosum* through the biphasic crystallization process. Other analytical grade chemicals were obtained from reputed manufacturers.

**Figure 1:**
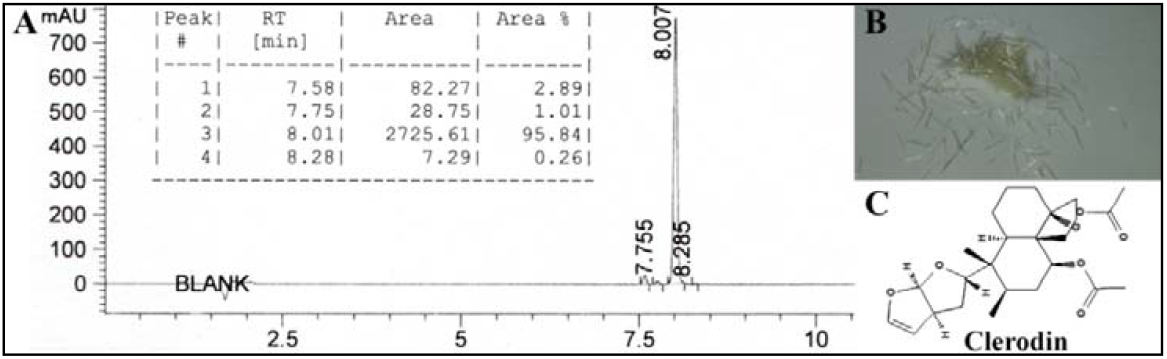
A; HPLC chromatogram and area percentage of clerodin. B; Isolated crystals of clerodin. C; 2-D conformer of clerodin (PubChem CIDs 442014).

### 2.2. Clerodin extraction from Clerodendrum viscosumVent

After collection of fresh *C. viscosum* leaves from Burdwan University campus, West Bengal, India, it was taxonomically identified, and a voucher specimen (No. BUTBSR011) is maintained in the Department of Zoology for future reference.

The collected fresh leaves were washed in tap water, dried in shade, ground by Philips Mixer Grinder HL1605, and the obtained leaf powder was stored in an airtight container for further use. 100 g of this pulverized leaf material was extracted in 2.5 L of boiling distilled water for 2-3 h and after that the extract was filtered with filter paper.

Leaf aqueous extract of *C. viscosum* (LAECV) was fractioned by petroleum ether with the help of a magnetic stirrer for 10-12 h and the resulting yellow colored petroleum ether extract was concentrated and dried by rotary vacuum evaporator and allowed for crystallization of clerodin for 48 h in hydro-methanolic solution. Needle shaped clerodin crystals were recrystallized to get 95.84% pure compounds (Figure 1).

### 2.3. Culture and treatment in human peripheral blood lymphocytes (HPBLs)

Human peripheral blood was collected with heparinized hypodermic syringe from healthy male donors. The 1 mL heparinized whole blood was cultured in sterilized 15 mL screw cap Tarson culture tubes using culture medium - RPMI 1640 supplemented with heat-inactivated FBS (10%), 2 mM L-Glutamine, and antibiotic solution [Penicillin-streptomycin (10,000 U/mL)]. Phytohaemagglutinine (PHA) was added to each culture tube as a mitogen. Culture tubes were incubated at 37°C in a CO_2_ incubator with 5% CO_2_ and humidified conditions within the incubator. HPBLs were allowed to grow for 36 h, after that of colcemid (0.1μg/mL) and 100μg/mL of clerodin was treated in respective culture tubes for 3 h and 6 h.

### 2.4. Culture and treatment in HEK293 cell line

HEK 293 (Human embryonic kidney293) cells were obtained from the National Centre for Cell Science (Pune, India). HEK cells were cultured in DMEM (high glucose) supplemented with 10% heat-inactivated FBS, Penicillin-streptomycin (10,000 U/ml) and 2 mM L-Glutamine in 25cm^2^ flasks that were incubated at 37°C in a CO_2_ incubator with 5% CO_2_ and humidified conditions within the incubator. 0.1μg/mL of colcemid and 100μg/mL of clerodin were treated in the respective culture tubes for 3 h and 6 h after 36 h of cell seeding. All the experiments were performed in triplicates.

### 2.5. Preparation of metaphase chromosomes

Peripheral blood lymphocytes were harvested by the conventional procedure (Ray and Chatterjee 2005), whereas, HEK 293 cells were harvested by 0.25% Trypsin-EDTA(1X) solution. HPBLs and HEK 293 cells were given hypotonic treatment (Hypotonic solution: 0.56% KCl prewarmed at 37°C) for 18 min and 22 min respectively and the cells were fixed in aceto-methanol (1:3) for 12 h. Clear, grease-free slides were used for the flame drying slide preparation method. The slides were stained with 5% Giemsa solution for 5-6 min and mounted in a synthetic medium.

### 2.6. MCF-7 cell line

#### 2.6.1. Cell cycle analysis by Flow cytometry

The flow cytometric cell cycle analysis was done using Propidium Iodide (PI), a fluorescent dye, binds to DNA stoichiometrically like a DNA-bound probe reflecting the cellular DNA content (Darzynkiewicz et al. 2004) and the frequency of cell cycle phases such as G1/G0, S, and G2/M-phase can be obtained by plotting the DNA fluorescence content in a histogram (Cunningham 1994).

The flow cytometric cell cycle analysis was done in MCF-7 cells after treating the cells with clerodin. Briefly, 1×10^6^ cells were seeded in 25cm^2^ flasks containing DMEM medium, supplemented with 10% FBS and antibiotic solution, and maintained at 37°C with 5% CO_2_. Upon 60% confluence, cells were treated with clerodin (0, 100, and 200μg/mL) for 24 h. The cells were fixed in 70% ethanol (chilled) for 30 min at 4°C. The fixed cells were washed in PBS to remove the ethanol followed by 50μg/mL RNase A was added to each tube and kept at room temperature for 1 h. Subsequently, 200μL Propidium Iodide (50 μg/mL) was added to each tube and kept at room temperature for 15-20 min. The samples were analyzed through the Beckman Coulter flow cytometer and FlowJo™ software (Treestar Inc, Ashland, OR) (Wang et al. 2019).

#### 2.6.2. Immunofluorescence imaging of tubulin by confocal microscope

Microtubule destabilization effects of clerodin were analyzed through confocal microscopic immunofluorescence imaging of tubulins, briefly,1×10^5^ MCF-7 cells were grown over lysine coated coverslips in 6-well sterile culture plates, upon 60% confluence, the cells were treated with clerodin (100 and 200μg/mL) for 24 h. The attached cells were fixed with 3.7% paraformaldehyde and subsequently washed in PBS. 0.1% Triton X-100 (v/v) in PBS for 10 min was used for cell permeabilization and blocking was done by 1% BSA in PBS for 2 h followed by washing in PBS. The cells were incubated with an anti-tubulin antibody overnight, washed in PBS, and subsequently incubated with FITC conjugated goat antimouse IgG for 2 h. Finally, the cells were washed in PBS 5 times (5 min each). After washing, the coverslips were mounted in DAPI/Antifade Solution. Simultaneously, similar preparation and processing of slides were done without incubation with the primary antibody, which regarded as a negative control. Prepared slides were observed and analyzed by a confocal laser scanning microscope (Carl Zeiss LSM 710, GmbH, Germany).

### 2.7. Root growth retardation and swelling effects of clerodin on *Allium cepa*

#### 2.7.1 Culture and treatment of Allium cepa roots

*Allium cepa* root apical meristem was used as a plant model for determining the root growth retardation inducing activities. 1% sodium hypochlorite mediated surface-sterilized *A. cepa* bulbs were placed in 6-well plates containing distilled water and kept in the environmental test chamber at 25-27 ° C for germination. Similar sized *A. cepa* roots with 2-3 cm root length (48 h aged) were treated with clerodin (0, 5, 10, 20, 40, and 80 μg/mL) continuously for 24, 48, and 72 h; simultaneously, colchicine was used as a positive control. Experiments were performed in triplicate.

### 2.8. Mitotic abnormality and polyploidy inducing effects of clerodin on *Allium cepa root tip cells*

#### 2.8.1. Treatment and preparation of mitotic phases

Microtubule destabilization based mitotic abnormalities and polyploidy inducing effects of clerodin and colchicine were analyzed on *A. cepa* root-tip cells. Colchicine was used here as a standard microtubule destabilizing compound which showed characteristic root swelling on *A. cepa* root tips in the recovery roots and induced different mitotic abnormalities, including, c-metaphase, micronuclei, and polyploidy. The similar-sized *A. cepa* roots (48 h aged) were treated with clerodin (0, 50, 100, and 150 μg/mL) and colchicine (150 μg/mL) for 2 to 4 h and then respectively 8-10 roots were processed for squash preparation. The remaining roots were allowed to grow further for another 16-32 h in distilled water and subsequently, the root tips were fixed in aceto-methanol (3 parts methanol: 1 part glacial acetic acid) for 24 h, hydrolyzed for 10 min in 1 N HCl at 60°C, stained with 2% aceto-orcein, and finally squashed in 45% acetic acid (Sharma and Sharma 1999, Ray et al. 2013). The well-spread areas of squashed roots were focused under the bright field light microscope (40X objective lens) for observation and scoring the cellular abnormality.

### 2.9. Clerodin induced polyploidy in radish (*Raphanus sativus*)

#### 2.9.1. Radish germination, treatment, and culture

Radish seeds were procured from local seed distributors and, approximately, 500 seeds were surface sterilized with 1% sodium hypochlorite and placed on sterile cotton filled Petri Dish containing distilled water. The Petri Dishes were kept at 23-25 ° C (12 h light: 12 h dark cycle) within an environmental test chamber for germination. Germinated seeds were placed in a 6-well plate culture disk for 24 h containing colchicine (50 μg/mL) and clerodin (30, 40 μg/mL) in the respective well. The untreated samples were maintained simultaneously in distilled water. At the end of 24 h of treatment, the seeds were gently rinsed with distilled water and carefully transferred to respective seedling pots containing fertile soil and continuously monitor for two months with a proper adequate water supply and to confirm the degree of polyploidy induced by clerodin and colchicine were analyzed on the basis of parameters like leaf length and breadth ratio, stomatal length, stomatal diameter, stomatal number, guard cell length, and finally, flow cytometric assessment of DNA contents in isolated protoplasts.

#### 2.9.2. Leaf length and breadth ratio, stomatal number, stomatal and guard cell size measurement

Record of leaf length and breadth ratio, stomatal number, stomatal and guard cell size are the suitable approach for polyploidy analysis (Omidbaigia et al.2010). For clerodin induced polyploidy assessment in radish, 6 plants from each treated and untreated groups were randomly selected for leaf length and breadth ratio determination. For stomatal and guard cell characteristics analysis, the nail varnish technique was performed. Clear nail polish was applied to the abaxial side of the leaf and allowed to dry for a few minutes. After drying, the layer of the nail polish was removed by fine tip forceps, placed on a grease-free slide, and observed under a bright-field microscope (LeitzLaborlux II, Germany) at 1000x magnification.

#### 2.9.3. Flow cytometric analysis for polyploidy

For detection of clerodin induced polyploidy, radish leaf protoplast DNA content was analyzed by flow cytometry. The protoplasts were isolated from the young radish leaves following the technique as described by Chawla H. (2009). Briefly, young leaves were first surface sterilized with 10% sodium hypochlorite containing 2 drops of Tween 20 for 10 min. With the help of fine forceps, the lower epidermis portion was peeled off from the leaves and then chopped into pieces with a scalpel. Leaf pieces were then transferred to 13% mannitol-CPW (cell and protoplast washing media) salt solution for 1 h. After that, the mannitol-CPW salt solution was replaced with 13% mannitol solution containing pectinase (0.5%) and cellulase (1%) for overnight. The enzyme mixture was then filtered and centrifuged to sediment the protoplast. The pellet was re-suspended into CPW salt solution containing 21% sucrose and again centrifuged at 100x g for 10 min and the uppermost band of the viable protoplasts was collected and washed with CPW salt solution containing 10% mannitol by centrifugation at 100x g for 10 min (3 times). The isolated protoplasts were fixed by 100% chilled methanol for 1 h. After fixation, methanol was removed and the protoplasts were incubated with RNase A (0.20mg/mL) at room temperature. Finally, stained with PI (Propidium Iodide, 0.04 mg/mL) for 15 min in the dark and the fluorescence intensity was recorded by CytoFlex, Beckman Coulter. Data were analyzed by FlowJo™ software (Treestar Inc, Ashland, OR).

### 2.10. Molecular docking analysis

The 3-D structures of the compounds colchicine and clerodin were obtained from the PubChem database (Bolton et al. 2008) bearing PubChem CIDs 6167 and 442014 respectively in the .sdf format. These 3-D structures were visualized using Molegro Molecular Viewer 2.5 and then converted to .mol2 format for carrying out docking studies (Berman et al. 2000). Ligands bound to the protein structure along with heteroatoms like water and ions were removed from the structure. This modified tubulin structure was carried forward for docking studies with colchicine and clerodin using Autodock 4.2 (Rizvi et al. 2013). The protein was loaded in the software followed by addition of polar hydrogen bonds and assigning Kollman charge to it. AutoGrid module (Goodford. 1985) was used to generate a grid box having dimensions of 126×106×82 along with a spacing angstrom of 0.819 to ensure that the grid box encompasses the whole protein. Lamarckian Genetic Algorithm was implemented to initiate docking experiments with an initial population size of 150, the gene mutation rate of 0.02, and a rate of cross-over of 0.8 (Rizvi et al. 2013). Ten independent docking runs were performed for each compound against tubulin. The binding energies of the protein-ligand complexes were calculated based on the equation:

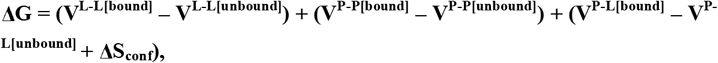

where; P refers to the protein; L refers to the ligand; V represents the pair-wise evaluations and S denotes the loss of conformational entropy upon binding.

Out of the ten independent docking runs, the conformation having the lowest binding energy represented the most favorable conformation and was visualized in UCSF Chimera 1.12 (Pettersen et al. 2004). Further, a more detailed molecular interaction study of the most favorable protein-ligand complexes were carried out using LIGPLOT (Wallace et al. 1995) to identify the protein residues involved in the interaction and their mode of interaction.

### 2.11. Scoring and statistical analysis

*Allium cepa* root growth retardation effect of clerodin was analyzed by student t-test. In the case of squash preparation of *A. cepa* root apical meristem cells, at least three randomly coded slides were observed under the light microscope. The mitotic index % was calculated by counting the number of dividing cells per total cells scored X 100 for each concentration. Aberrant cell percentage was calculated by counting the number of abnormal cells scored per total cells scored X 100 cells for each concentration (Bakare et al. 2000). Different cell phase frequencies, mitotic index, and mitotic abnormalities were analyzed by 2X2 contingency χ_2_-test.

## 3. Results

### 3.1. Metaphase arrest and c-metaphase analysis in HPBLs

Mitotic index (MI; metaphase frequency) was analyzed in HPBLs following treatment with clerodin (100 μg/mL) or colcemid (0.1 μg/mL) for 3 and 6 h. The average frequency of mitotic index in untreated control was 0.26±0.03% and 0.40±0.02% at 39 and 42 h of culture, respectively. The frequency of MI was increased to 1.76±0.18 and 2.88±0.19% after treatment with clerodin for 3 and 6 h, respectively. Following treatment with colcemid for 3 and 6 h, the average frequency of MI was 3.51±0.18% and 4.59±0.16%, respectively (Figure 2, 4 and Table **S**1).

**Figure 2:**
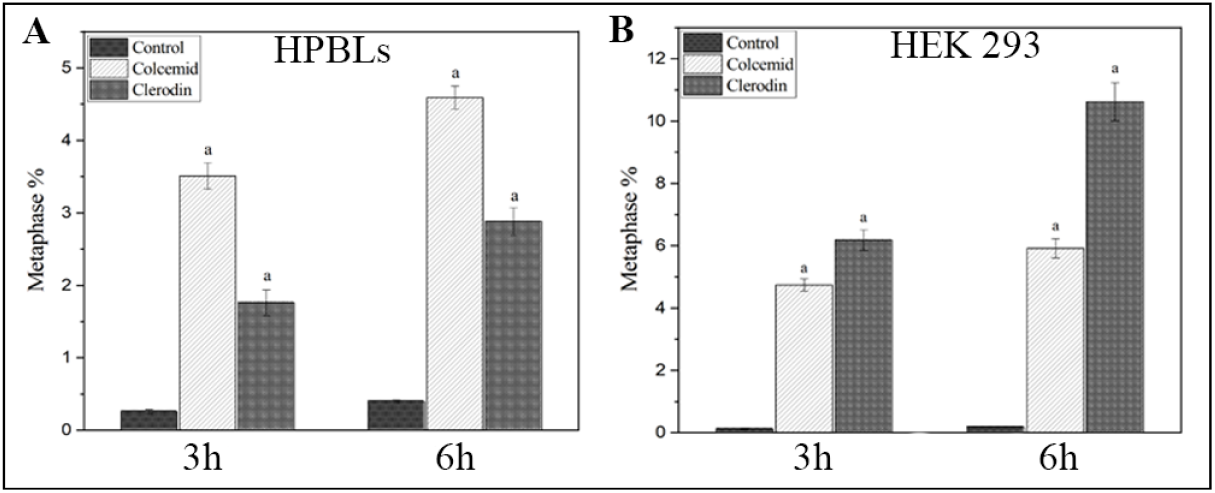
Effects of clerodin and colcemid on the metaphase percentage in human peripheral blood lymphocyte **(A)** and HEK 293 cells (B). Data represented as Mean±SEM. ^a^ significant at p<0.001 as compared to their respective control by 2×2 Contingency χ^2^-test with respective d.f. = 1.

**Figure 3.**
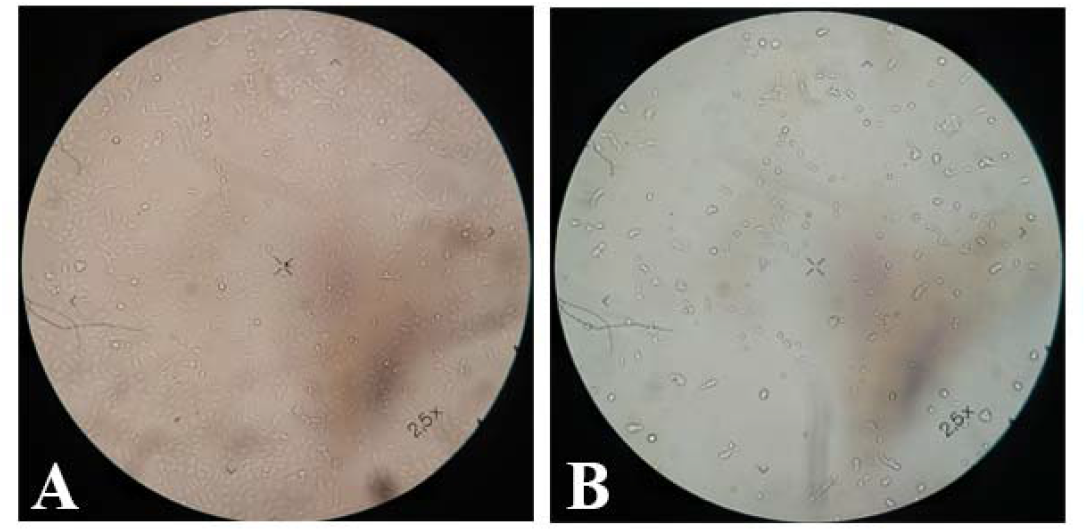
HEK 293 cell culture plates showing growth inhibitory effects of clerodin. A. untreated and B. clerodin treated.

**Figure 4.**
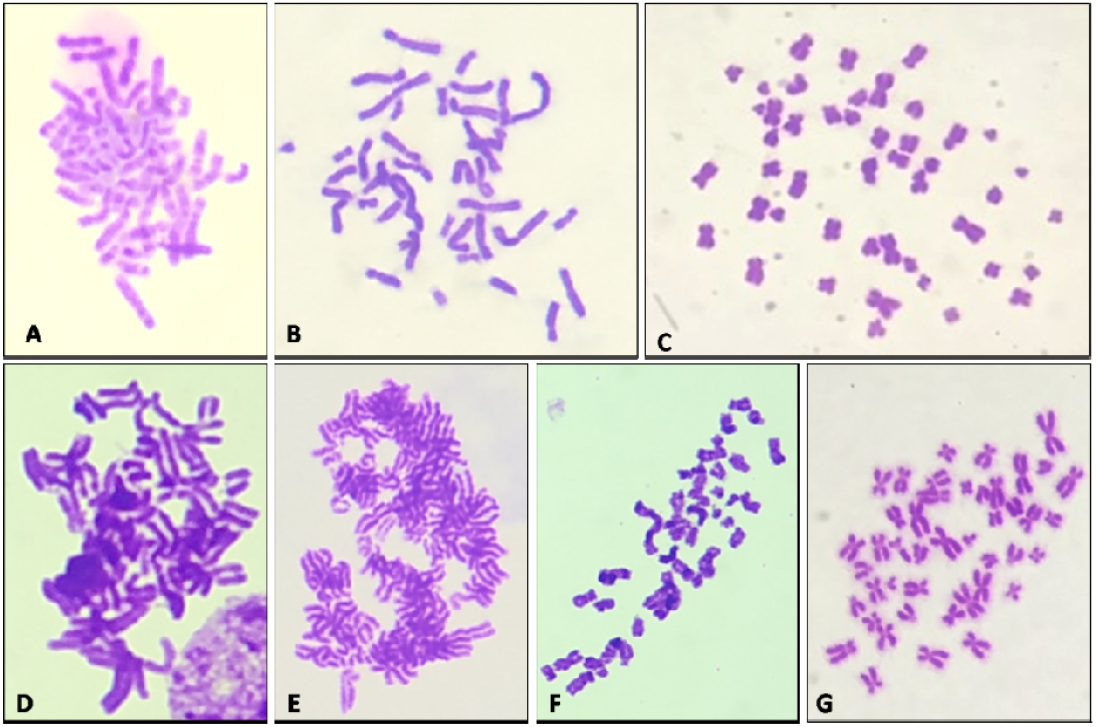
Clerodin and colcemid induced metaphase chromosome plates of HPBLs (A, B, C) and HEK 293 cells (D, E, F, G). A; D; Untreated cells, B; E; F; 100μg/mL of clerodin and C; G; 100μg/mL treated colcemid.

### 3.2. Metaphase arrest and c-metaphase analysis in HEK 293 cells

Like HPBLs, in HEK 293 cell line the standard drug colcemid induced 4.74±0.2 and 5.92±0.31% MI while clerodin (100μg/mL) induced 6.18±0.33 and 10.62±0.61% at 3 and 6 h of treatment, respectively (Figure 2, 4).

### 3.3. Flowcytometry and confocal microscopic analysis of MCF-7 cells

Data indicate that clerodin has the potential to increase G2-M frequencies in the MCF-7 cell line. 100 and 200μg/mL concentration of clerodin induced 11.28 and 16.25% G2-M frequencies, whereas 6.10% G2-M frequencies were recorded in untreated cells. G1 and S phase frequencies show a sharp decrease pattern in clerodin treatment compared to untreated. Here, G1 frequencies recorded 34.32% from the untreated cells and that decreased to 21.62% (100 μg/mL) and 21.28% (200μg/mL) at 24 h clerodin treatment. Similarly, 55.88% of S phase frequencies were obtained in the untreated cells, which reduced to 50.51% at 200μg/mL of clerodin treatment (Figure 5, A-C).

**Figure 5:**
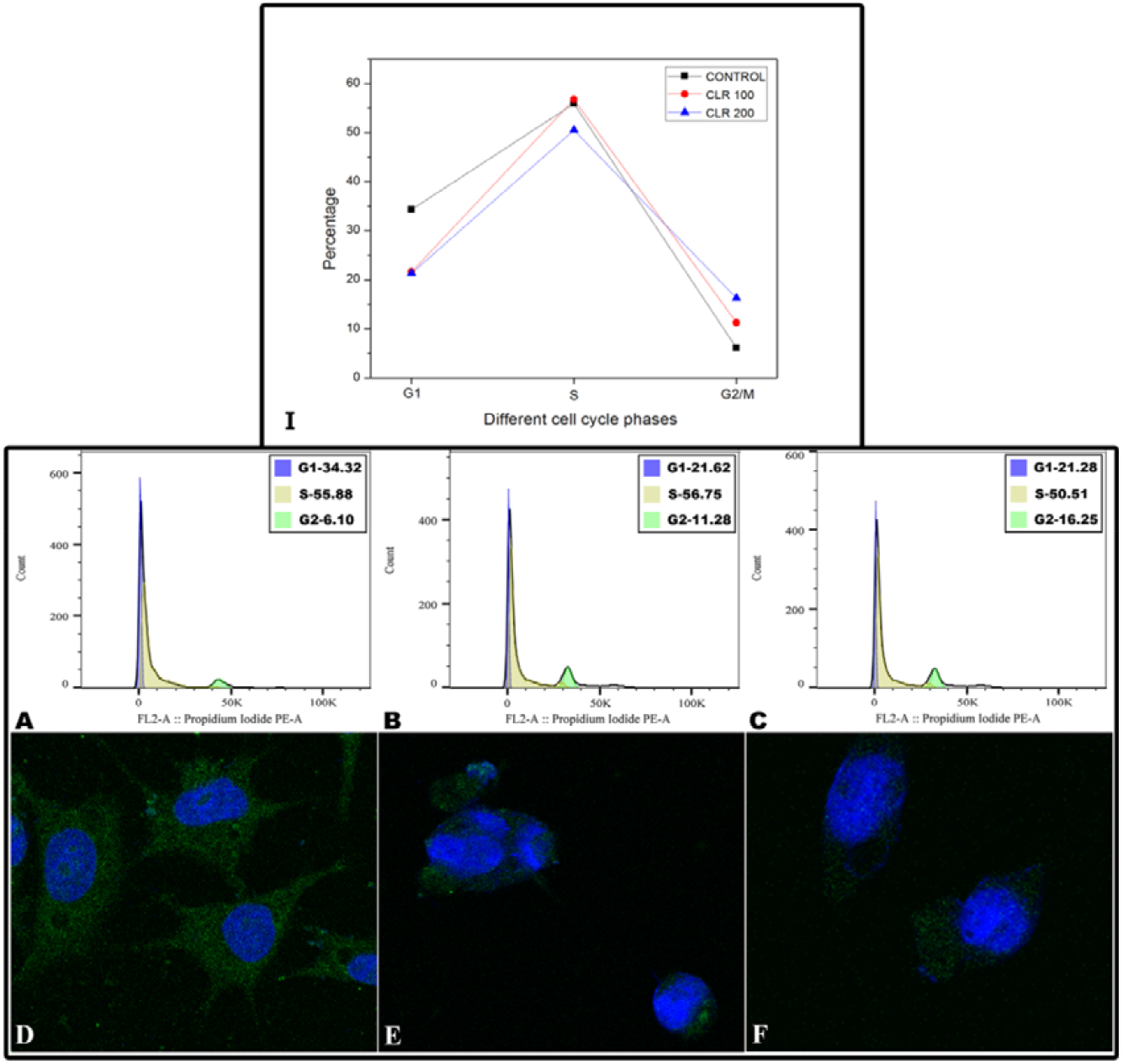
Effect of clerodin on cell cycle phase-frequencies and immunofluorescence imaging of tubulin in MCF-7 cells. I; Different cell cycle phase-frequency percentage, A; D; Untreated cells, B; E; 100μg/mL and C; F; 200μg/mL of clerodin treated in MCF-7 cell line. Tubulin and nuclei were visualized in green and blue respectively.

The confocal microscopic imaging of tubulin in MCF-7 cells showed that clerodin reduced the tubulin network and the microtubule depolymerization may be the basis of clerodin induced G2-M arrest in MCF-7 cells (Figure 5, D-F).

### 3.4. Apical meristem

#### 3.4.1. Allium cepa root growth retardation and root swelling

Data indicate both clerodin and colchicine treatment-induced time (24-72 h) and concentration-dependent (10-80 μg/mL) growth retardation on *A. cepa* roots (Figure 6; A & Table **S**2). Root growth retardation was recorded at 90.42±1.83 and 84.08±1.41% respectively in continuous treatment of clerodin and colchicine treatment (80 μg/mL) for 72 h. The maximum root growth retardation (58.28±1.24%; p<0.001 at 72h) was recorded with the highest concentration (80 μg/mL) of clerodin treatment while at the same concentration of colchicine the growth retardation (50.32±1.52%; p<0.001) was comparatively lower. The root growth retardation IC_50_ values for clerodin and colchicine were respectively 18.98±4.16 and 29.83±2.12 μg/mL (Figure 6; B).

**Figure 6:**
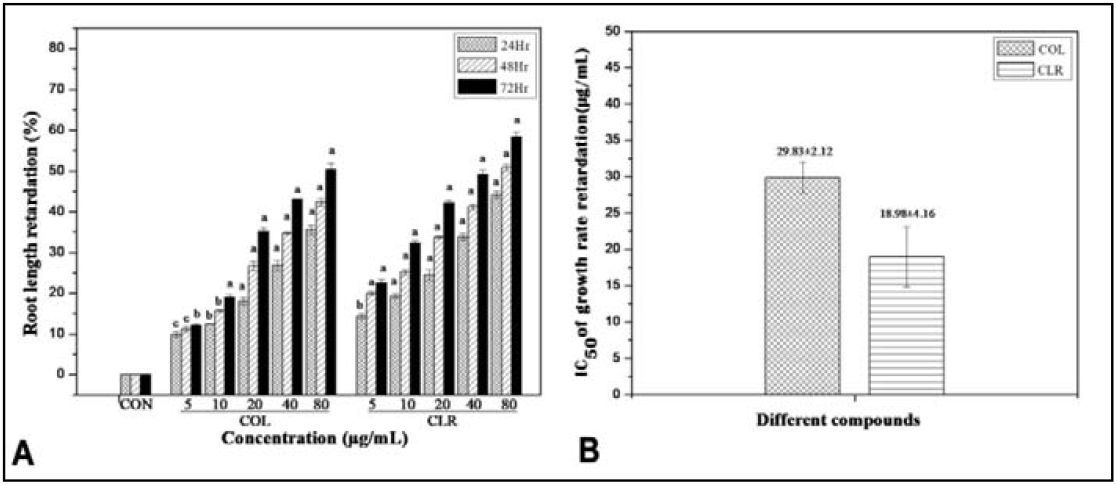
Root length retardation effect of clerodin and colchicine (A) and IC_50_ of the root growth retardation (B) on *Allium cepa*. Data represented as Mean ± SEM. ^a^ significant at *p*<0.001 and ^b^ significant at *p*<0.01, ^c^ significant at *p*<0.05 as compared to their respective control by Student’s t-test with respective d.f. = 1.

#### 3.4.2. Alteration of the mitotic index and mitotic phase-frequency

The mitotic index of untreated root apical meristem cells were 5.99 ± 0.04, 7.13 ± 0.13, 8.95 ± 0.05, and 8.51 ± 0.37% respectively at 2, 4, 20, and 36 h. The mitotic index percentage increased with the increasing concentration of clerodin at 2 h (12.9±0.17% at150 μg/mL) and 4 h (13.44±0.57% at 100 μg/mL) treatment, but decreases at 4+16 h (7.77±0.21 at 150 μg/mL; p<0.05) and 4+32 h (5.04±0.16 at 50 μg/mL) treatment. Colchicine also showed similar mitotic index modulatory effects. Dose-dependent reduction of prophase, anaphase, and telophase frequency was induced by clerodin and colchicine but metaphase frequency increases in the case of both treatments (Table 1& Figure 7).

**Figure 7:**
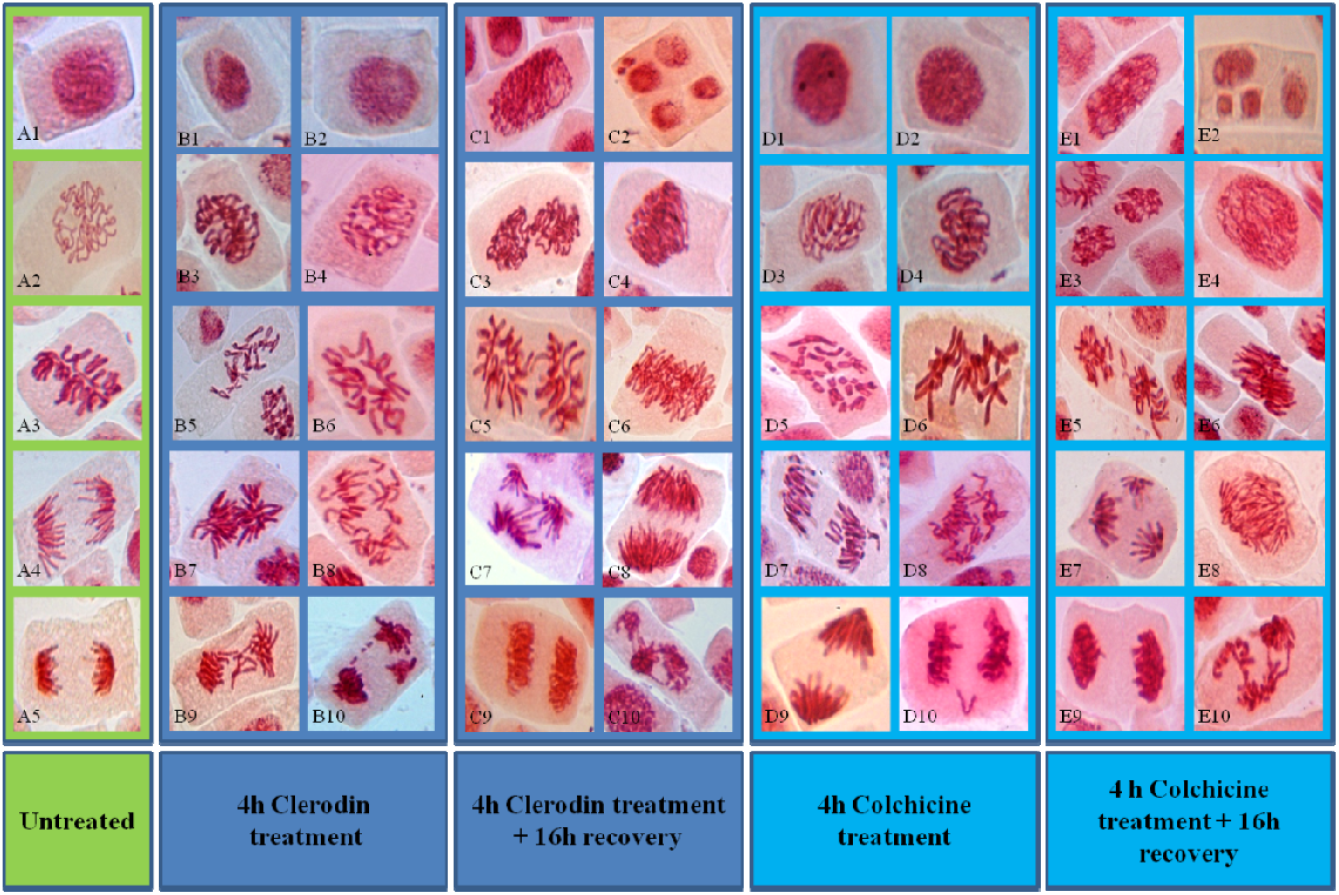
Different cell division phases and mitotic abnormalities induced by clerodin and colchicine in *A. cepa* root apical meristem cells. A1-A5; Untreated Interphase, Prophase, Metaphase, Anaphase, and Telophase. B1-B4; Interphase and Prophase from Clerodin treatment for 4 h. B5-B6; Different mitotic abnormalities induced by Clerodin at 4 h treatment. C1-C10; Different cellular and mitotic abnormalities induced by Clerodin at 4 h Clerodin + 16 h recovery. D1-D4; Interphase and Prophase from Colchicine treatment for 4 h. D5-D10; Different mitotic abnormalities induced by Colchicine at 4 h treatment. E1-E10; Different cellular and mitotic abnormalities induced by Colchicine at 4 h Colchicine treatment +16h recovery. A1, B1, B2, D1, D2; Interphase. A2, B3, B4, D3, D4; Prophase. A3; Metaphase. A4; Anaphase. A5; Telophase. B5, D5; C-metaphase. B6, D6; Disrupted metaphase. B7, D9; Polar deviation. B8, D8; Disrupted anaphase. B9, D7; Laggard chromosome. B10, D10; Sticky and Vagrant chromosome. C1, E1; Polyploid interphase. C2, E2; Multiple nuclei with abnormal cell division furrow and Micronucleus. C3, E3; Binucleate prophase. C4, E4; Polyploid prophase. C5, E5; Binucleate metaphase. C6, E6; Polyploid metaphase. C7, E7; Multipolar anaphase. C8, E8; Polyploid anaphase. C9, E9; Polyploid telophase. C10, E10; Multipolar telophase with bridge.

**Table 1:** Effects of clerodin and colchicine on the mitotic phase frequency in *A. cepa* root apical meristem cells.

| H | Conc<br>(ug/mL) | TC | DC | MI(%) | Pro(%) | Meta(%) | Ana(%) | Telo(%) |
| --- | --- | --- | --- | --- | --- | --- | --- | --- |
| **2** | 0 | 5572 | 334 | 5.99±0.04 | 20.4±1.17 | 34.34±1.2 | 27.89±0.84 | 17.33±0.68 |
|  | 50 | 6518 | 492 | 7.53±0.14<sup>b</sup> | 15.60±0.39 | 56.65±0.36 | 19.59±0.64 | 8.16±0.16 |
|  | 100 | 6779 | 758 | 11.20±0.40<sup>a</sup> | 16.89±0.36 | 62.65±0.49 | 14.9±0.39 | 5.55±0.44 |
|  | 150 | 7673 | 992 | 12.9±0.17<sup>a</sup> | 19.55±0.28 | 64.92±0.13 | 9.28±0.42 | 6.25±0.24 |
|  | 150© | 7224 | 851 | 11.77±0.04<sup>a</sup> | 10.56±0.25 | 76.59±0.42 | 8.12±0.26 | 4.73±0.46 |
| **4** | 0 | 8102 | 578 | 7.13±0.13 | 20.03±0.78 | 35.68±0.76 | 30.79±0.64 | 13.49±0.28 |
|  | 50 | 8339 | 1120 | 13.4±0.26<sup>a</sup> | 10.79±0.13 | 66.82±0.66 | 16.48±0.47 | 5.91±0.21 |
|  | 100 | 11381 | 1527 | 13.44±0.57<sup>a</sup> | 9.22±0.24 | 77.42±0.38 | 9.17±0.14 | 4.19±0.15 |
|  | 150 | 13339 | 1738 | 13.05±0.29<sup>a</sup> | 9.6±0.18 | 78.21±0.68 | 8.56±0.41 | 3.62±0.35 |
|  | 150© | 7292 | 655 | 8.98±0.05<sup>a</sup> | 8.10±0.23 | 79.42±1.04 | 7.01±0.26 | 5.46±0.80 |
| **4+16** | 0 | 5806 | 520 | 8.95±0.05 | 22.11±0.22 | 36.15±0.16 | 27.88±0.19 | 13.85±0.44 |
|  | 50 | 8482 | 432 | 5.08±0.06<sup>a</sup> | 15.97±0.18 | 47.89±0.29 | 24.36±0.38 | 11.79±0.15 |
|  | 100 | 5834 | 392 | 6.69±0.30<sup>a</sup> | 10.69±0.14 | 51.32±0.57 | 26.94±1.33 | 11.04±0.82 |
|  | 150 | 4666 | 363 | 7.77±0.21<sup>c</sup> | 8.27±0.27 | 59.02±0.99 | 24.22±0.6 | 8.48±0.6 |
|  | 150© | 5495 | 187 | 3.37±0.31<sup>a</sup> | 21.73±1.06 | 41.91±0.77 | 16.21±0.74 | 20.14±0.7 |
| **4+32** | 0 | 7226 | 613 | 8.51±0.37 | 31.88±0.99 | 33.05±1.43 | 21.12±1.33 | 13.93±0.79 |
|  | 50 | 8297 | 418 | 5.04±0.16<sup>a</sup> | 30.61±1.35 | 38.53±1.65 | 21.75±0.81 | 9.09±0.32<sup>c</sup> |
|  | 100 | 8789 | 375 | 4.26±0.18<sup>a</sup> | 22.77±1.66<sup>c</sup> | 46.94±0.92<sup>c</sup> | 22.46±1.29 | 7.81±0.98<sup>c</sup> |
|  | 150 | 9478 | 404 | 4.26±0.15<sup>a</sup> | 21.54±0.50<sup>c</sup> | 48.19±0.90<sup>b</sup> | 20.07±0.87 | 10.20±0.54 |
|  | 150© | 9029 | 424 | 4.67±0.27<sup>a</sup> | 34.01±0.80 | 43.82±1.07<sup>c</sup> | 12.99±1.64<sup>c</sup> | 9.30±0.53<sup>c</sup> |

©=Colchicine treated, <sup>a</sup> significant at p<0.0001 and <sup>b</sup> significant at p<0.001, <sup>c</sup> significant at p<0.05 as compared to their respective control by 2x2 Contingency $\chi^2$ -test with respective d.f.= 1. H; Hours, Comp; Compound, Conc; Concentration, TC; Total no of cells, DC; No of dividing cells, MI; Mitotic index, Pro; Prophase, Meta; Metaphase, Ana; Anaphase, Telo; Telophase.

#### 3.4.3. Aberrant cell percentage

Both clerodin and colchicine treatment could increase aberrant cell frequency in onion root tip cells in a dose-dependent manner at 2 and 4 h. The maximum aberrant cell percentage (10.33 ± 0.3 and 11.33 ± 0.4) was recorded at 150 μg/mL concentration of clerodin whereas the same used concentration of colchicine could induce a similar trend of higher aberrant cells percentage (9.10 ± 0.21 and 7.6 9± 0.15% respectively at 2 and 4 h). In the case of 16 h recovery samples, the aberrant cell frequency reduced in both clerodin (5.23±0.22 at 150 μg/mL) and colchicine (0.65±0.05 at150 μg/mL) (Table 2& Figure 7).

**Table 2:** Effects of clerodin and colchicine on the frequency of mitotic abnormalities induced in *A. cepa* root apical meristem cells.

| H | Conc (ug/mL) | Tac<br>(%) | Sti<br>(%) | Bri<br>(%) | Pd<br>(%) | C Met<br>(%) | Vag<br>(%) | Lag<br>(%) |
| --- | --- | --- | --- | --- | --- | --- | --- | --- |
| 2 | 0 | 0.2±0.02 | 2.09±0.35 | 1.19±0.30 | 0±0 | 0±0 | 0±0 | 0±0 |
|  | 50 | 4.89±0.10<sup>a</sup> | 13.08±0.50<sup>a</sup> | 11.78±0.27<sup>a</sup> | 5.06±0.12<sup>a</sup> | 46.25±0.55<sup>a</sup> | 1.64±0.25 | 1.41±0.09 |
|  | 100 | 8.15±0.4<sup>a</sup> | 13.18±0.32<sup>a</sup> | 12.23±0.51<sup>a</sup> | 8.04±0.37<sup>a</sup> | 53.2±1.11<sup>a</sup> | 3.55±0.11<sup>b</sup> | 2.09±0.28 |
|  | 150 | 10.33±0.3<sup>a</sup> | 15.73±0.69<sup>a</sup> | 7.36±0.56<sup>a</sup> | 6.04±0.43<sup>a</sup> | 61.68±0.63<sup>a</sup> | 4.03±0.55<sup>b</sup> | 1.71±0.36 |
|  | 150© | 9.10±0.21<sup>a</sup> | 2.22±0.25 | 3.41±0.31<sup>c</sup> | 1.17±0.09 | 70.47±0.28<sup>a</sup> | 0.70±0.20 | 0±0 |
| 4 | 0 | 0.09±0.01 | 1.39±0.20 | 0±0 | 0±0 | 0±0 | 0±0 | 0±0 |
|  | 50 | 9.71±0.08<sup>a</sup> | 13.92±0.07<sup>a</sup> | 14.14±0.5<sup>a</sup> | 7.15±0.18<sup>a</sup> | 52.03±0.97<sup>a</sup> | 2.78±0.11<sup>a</sup> | 1.19±0.31 |
|  | 100 | 10.72±0.38<sup>a</sup> | 14.15±0.33<sup>a</sup> | 7.66±0.22<sup>a</sup> | 7.14±0.07<sup>a</sup> | 74.11±0.50<sup>a</sup> | 3.59±0.08<sup>a</sup> | 1.37±0.08<sup>c</sup> |
|  | 150 | 11.33±0.4<sup>a</sup> | 15.24±0.64<sup>a</sup> | 7.58±0.17<sup>a</sup> | 7.48±0.12<sup>a</sup> | 74.62±0.18<sup>a</sup> | 3.68±0.20<sup>a</sup> | 1.44±0.06<sup>c</sup> |
|  | 150© | 7.69±0.15<sup>a</sup> | 0.92±0.02 | 6.11±0.43<sup>a</sup> | 3.06±0.34<sup>a</sup> | 74.1±1.22<sup>a</sup> | 0.77±0.17 | 0±0 |
| 4+16 | 0 | 0.15±0.003 | 1.73±0.02 | 0±0 | 0±0 | 0±0 | 0±0 | 0±0 |
|  | 50 | 2.08±0.04<sup>a</sup> | 3.0±0.09 | 6.79±1.02<sup>a</sup> | 6.45±0.20<sup>a</sup> | 25.99±1.66<sup>a</sup> | 1.65±0.33 | 0.95±0.29 |
|  | 100 | 3.84±0.06<sup>a</sup> | 6.73±1.07<sup>b</sup> | 10.69±0.74<sup>a</sup> | 8.69±0.54<sup>a</sup> | 36.66±1.06<sup>a</sup> | 1.29±0.28 | 0.54±0.27 |
|  | 150 | 5.23±0.22<sup>a</sup> | 8.05±0.62<sup>a</sup> | 12.32±0.90<sup>a</sup> | 12.32±0.68<sup>a</sup> | 46.27±2.8<sup>a</sup> | 3.01±0.13 | 1.09±0.25 |
|  | 150© | 0.65±0.05<sup>a</sup> | 0±0 | 2.08±0.23 | 3.3±0.4 | 11.22±0.37<sup>a</sup> | 2.66±0.45 | 0±0 |

©=Colchicine treated, <sup>a</sup> significant at p<0.0001 and <sup>b</sup> significant at p<0.001, <sup>c</sup> significant at p<0.05 as compared to their respective control by 2x2 Contingency $\chi^2$ -test with respective d.f. = 1. H; Hours, Comp; Compound, Conc; Concentration, Tc; Total number of cells, TDC; Total dividing cells, Ac; Aberrant cells, Sti; Stickiness, Bri; Bridge, Pd; Polar deviation, C-Meta; Colchicine like Metaphase, Vag; vagrant chromosome, Lag; Laggard chromosome.

#### 3.4.4. Microtubule destabilization associated mitotic abnormalities in onion root tip cells

The different types of mitotic abnormalities like c-metaphase, polar deviation, vagrant chromosome, lagging chromosome, and micronucleus that are associated with the microtubule destabilization induced by clerodin and colchicine treatment are scored from squashed *Allium cepa* root apical meristem cells.

#### 3.4.5. C-metaphase frequency

Clerodin and colchicine treatment for 2 and 4 h increased the C-metaphase frequency in a dose-dependent manner and its highest frequency was observed at 4h treatment. Clerodin and colchicine (150 μg/mL concentration) induces 74.62 ± 0.18 and 74.1 ± 1.22% of C-metaphase respectively at 4h treatment. After 16 h of recovery treatment, the induced C-metaphase frequency 46.27±2.8% at 150 μg/mL and 11.22±0.37% at 150 μg/mL comparatively reduced respectively in clerodin and colchicine (Table 2& Figure 7, 10).

#### 3.4.6. Polar deviation

Data indicate that both clerodin and colchicine treatments could induce an increased percentage of polar deviation in the treated onion root tip cells. The highest polar deviation frequency (12.32±0.68%) was observed at 4+16 h recovery treatment of clerodin (150μg/mL) and it was far greater than colchicine (3.3±0.4%) affects on *Allium cepa* root apical meristem cells (Table 2& Figure 7).

#### 3.4.7. Vagrant chromosome

The data indicate that the used highest concentration (150μg/mL) of clerodin treatment induced higher frequencies (4.03±0.55% and 3.01±0.13%) of the vagrant chromosome at early hours (2 and 4 h) of continuous treatment, whereas, in case of 4 h treatment followed by 16 h recovery samples, the overall vagrant chromosome frequency reduced but increased the percentage (3.01±0.13 at150 μg/mL) of cells with vagrant chromosomes with the increasing concentrations of clerodin (Table 2& Figure 7).

#### 3.4.8. Laggard chromosome

Like vagrant chromosome, the increased percentage of cells (2.09±0.28 %) with laggard chromosome can be seen in clerodin treatment (100μg/mL) for 2 h (Table 2 & Figure 7). The frequency of laggard chromosome decreased at 4 h of treatment and also in16 h recovery samples.

#### 3.4.9. Anaphase bridge

Data indicate clerodin treatment could induce an increased percentage of anaphase bridges. The highest frequency of anaphase-bridge (14.14±0.5%) was observed at a concentration of 50 μg/mL of clerodin at 4 h of treatment but with the increasing dose of clerodin the frequency of anaphase reduced. Colchicine (150 μg/mL) treatment for 2 and 4 h respectively induced anaphase bridge frequency of 3.41 ± 0.31% and 6.11 ± 0.43%. In the case of 4 h treatment followed by 16 h recovery samples, the anaphase bridge frequency reduced in colchicine (2.08±0.23%) treatment but there was a dose-dependent increase (12.32±0.90%) in clerodin (150 μg/mL) treated samples (Table 2& Figure 7).

#### 3.4.10. Chromosomal stickiness

Data indicate that both clerodin and colchicine could induce chromosomal stickiness in root apical meristem cells and the frequency of stickiness increases in a dose-dependent manner. Clerodin treatment (150μg/mL) for 2 and 4 h respectively show sticky chromosome frequency of 15.73±0.69% and 15.24±0.64%.Colchicine induces sticky chromosome at 2 and 4 h treatment only whereas clerodin also induces increased sticky chromosome frequency at 4 h treatment followed by16 h recovery (Table 2 & Figure 7).

#### 3.4.11. Micronucleus

In the case of 16 h recovery treatments, both clerodin (50μg/mL) and colchicines (150μg/mL) induced increased micronucleus frequency respectively as 4.39±0.12 and 7.29±0.35% on *A. cepa* root apical meristem cells. The induced micronucleus frequency increased (7.97±0.38 %) at 32 h recovery samples after clerodin (150μg/mL) treatment but it is slightly reduced (6.71±0.26%) in the colchicine (150μg/mL) treated samples (Figure 7, 8 & 10).

**Figure 8:**
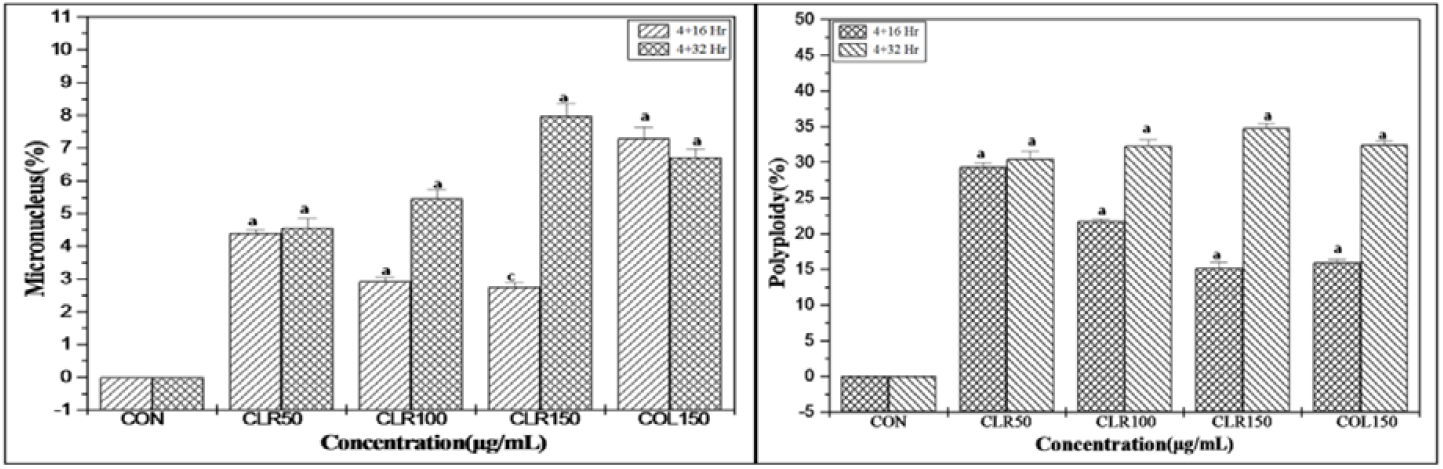
Effects of clerodin and colchicine on the frequency of micronuclei and polyploidy induced in *A. cepa* root apical meristem cells. Data represented as Mean±SEM. ^a^ significant at p<0.0001 and ^b^ significant at p<0.001, ^c^ significant at p<0.05 as compared to their respective control by 2×2 Contingency χ^2^-test with respective d.f. = 1. CON; Control, CLR; Clerodin, COL; Colchicine.

#### 3.4.12. Polyploidy

In the case of both clerodin (50 μg/mL) and colchicine (150 μg/mL) treatment (4 h) followed by 16 h recovery samples, the increased polyploid cells frequencies were 29.35±0.50 and 15.95±0.42% respectively in root apical meristem cells. The polyploid cell frequency increased further at 32 h recovery samples where clerodin and colchicine treatments (150 μg/mL) induced 34.80±0.62 and 32.45±0.56% of polyploidy cells frequency respectively (Figure 7, 8, 9, 10 & **S**10).

**Figure 9:**
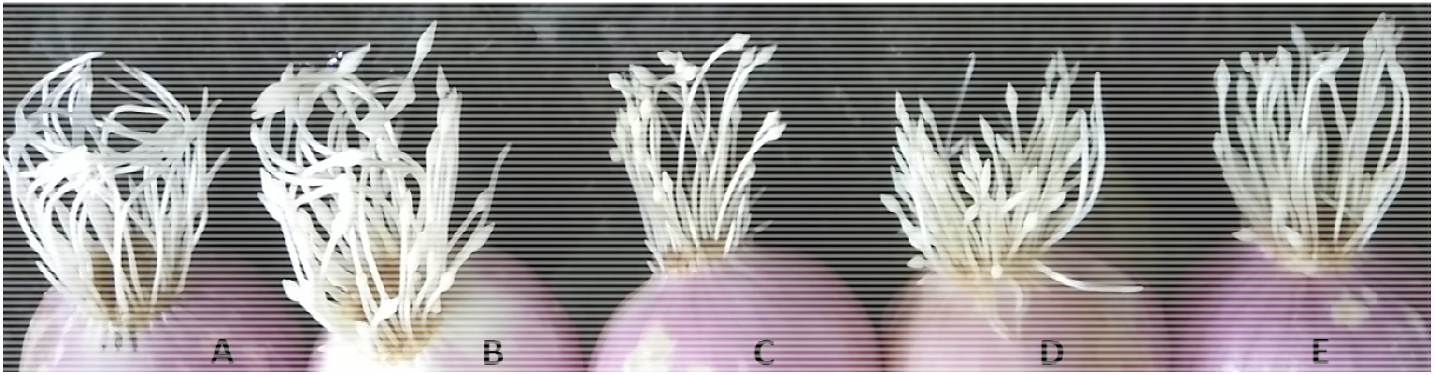
Clerodin and colchicine induced root tip swelling pattern of *A. cepa* at 4 h treatment followed by 32 h recovery. A; Control, B; Colchicine (150 μg/mL), C; Clerodin(50 μg/mL), D; Clerodin (100 μg/mL), E; Clerodin (150 μg/mL).

**Figure 10:**
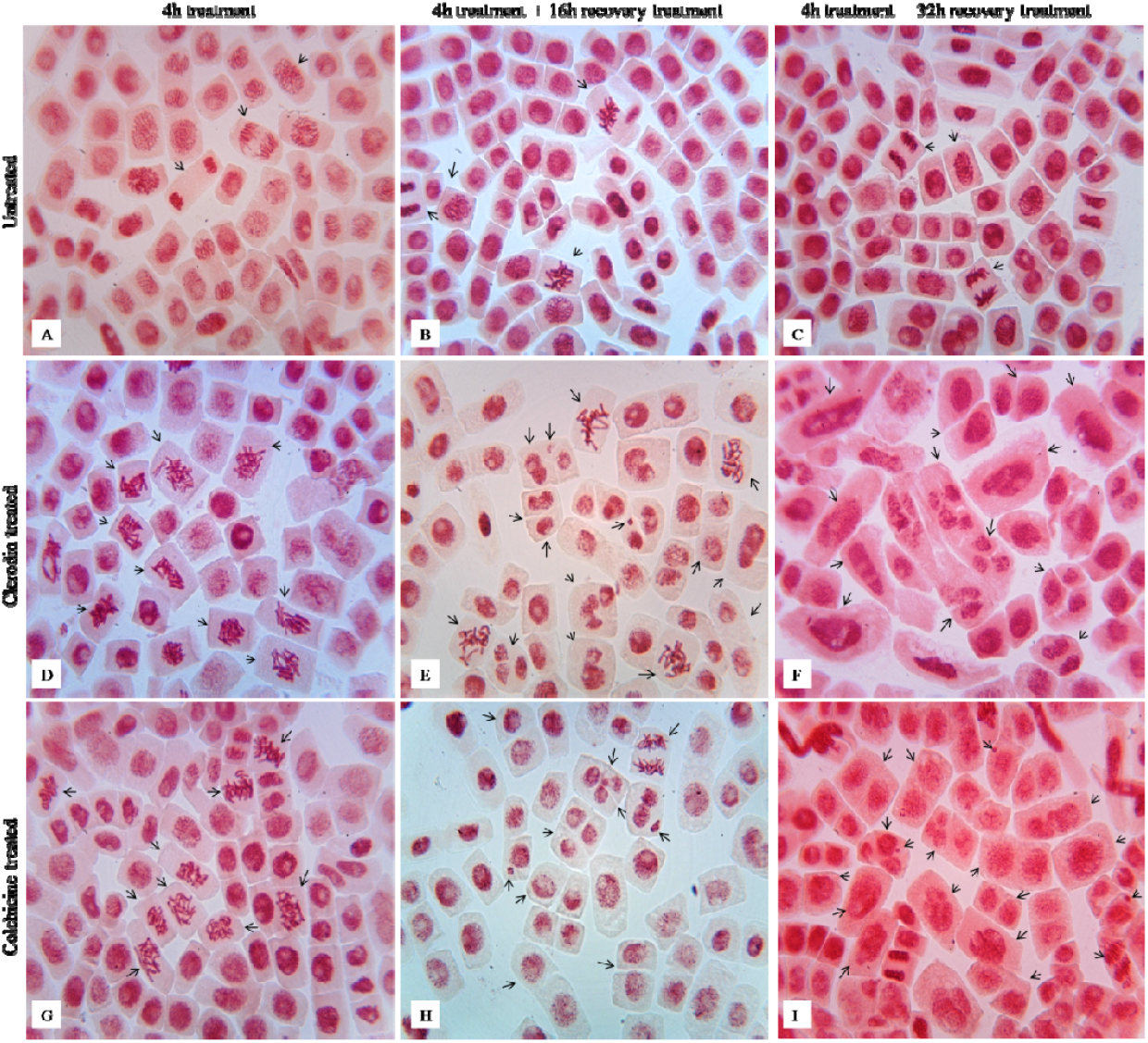
Photomicrographs showing squashed root tips; the C-metaphase, Polyploidy and Micronucleus frequency induced by Clerodin and Colchicine in *A. cepa* root apical meristem cells at 4 h, 4+16 h, and 4+32 h treatment. A, B, C; Control plates. D, E, F; Clerodin treated plates. G, H, I; Colchicine treated plates.

**Figure S10.**
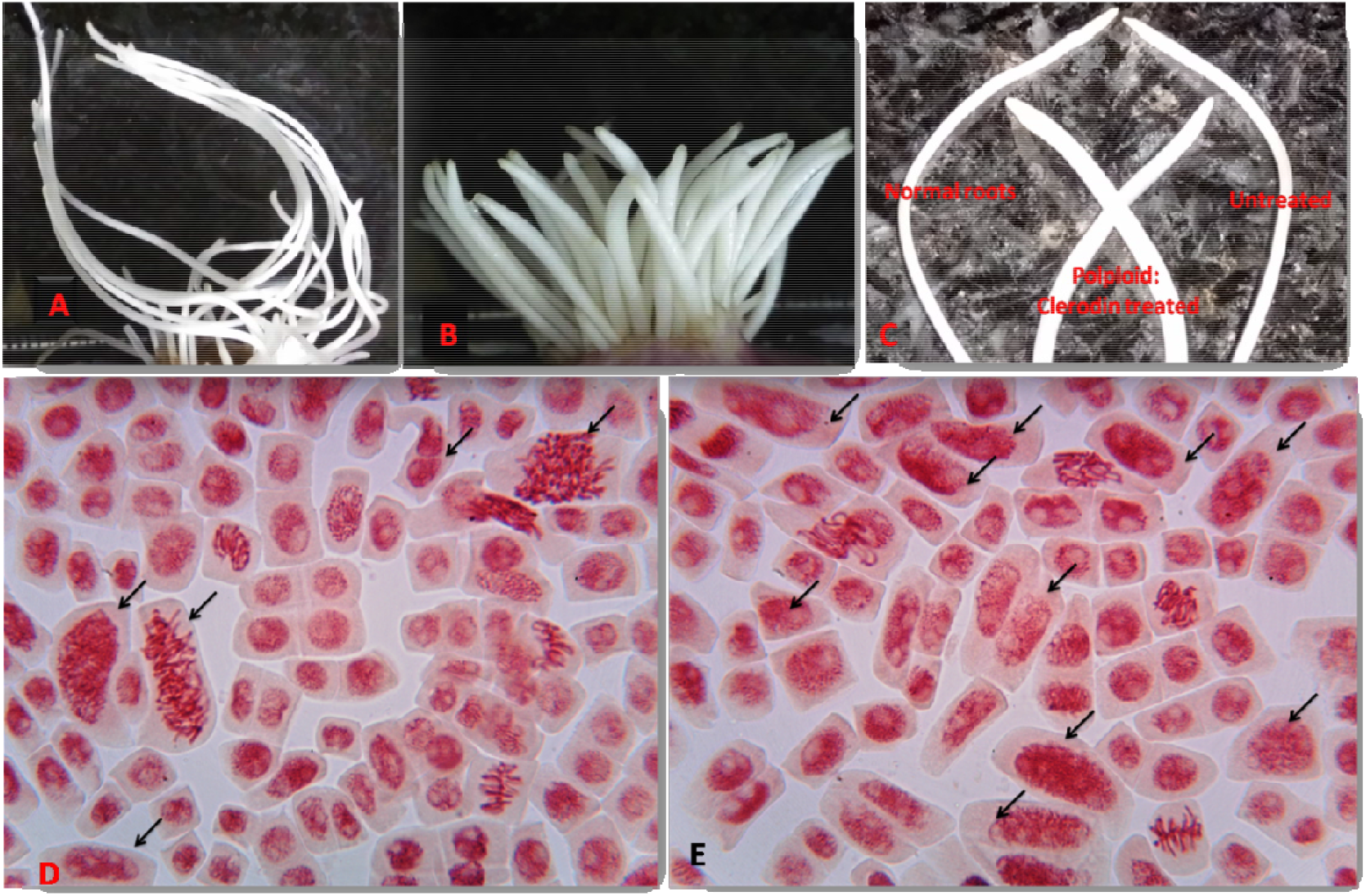
Clerodin induced polyploid roots. A; Untreated, B; Clerodin induced polyploid roots, C; showing comparison between normal and polyploid roots, D & E; clerodin treated squashed roots showing induced polyploid cells.

### 3.5. Radish polyploidy

#### 3.5.1. Leaf morphometric analysis

Data indicate that both clerodin and colchicine treatment-induced polyploidy in radish. Measurement of leaf length/breadth ratio, stomatal number, stomatal and guard cell size are considered as an indicator of induced polyploidy. The length of the stomata increased in colchicine (18.42±0.23 μm at 50 μg/mL) and clerodin (23.33±0.36 μm at 40 μg/mL) treated samples as compared to untreated (11.18±0.10 μm) (Figure 11 & Table **S**3). Like stomatal length, the stomatal diameter also increased in treated samples. In comparison to untreated (1.53±0.03), 50 μg/mL of colchicine and 30, 40 μg/mL of clerodin treatment resulted in the stomatal diameter of 2.78±0.05, 3.65±0.07, and 3.83±0.08 μm respectively. An increase in stomatal length and diameter is also related to a decrease in stomatal number present in per square millimeter leaf surface. Depiction of stomatal number by nail varnish technique indicate that about half fold reduction in colchicine (5.07±0.14 at 50 μg/mL) and clerodin (4.33±0.19 at 40 μg/mL) treatment compared to untreated (13.41±0.22). Guard cell length also tended to grow with stomatal length in colchicine (26.56±0.37 μm at 50 μg/mL) and clerodin (33.60±0.39 μm at 40 μg/mL) treatment compared to untreated (16.56±0.14) (Figure 11). Both clerodin and colchicine treatments altered the leaf length and breadth ratio in radish plants, and the ratio decreased in treated samples as compared to untreated controls. A decrease in length and breadth ratio indicates that the leaves became shorter and wider in shape, which is a characteristic feature of polyploid plants. In the present study, the highest leaf length and breadth ratio scored in case of untreated (2.02±0.02) samples and that reduced in both colchicine (1.40±0.02 at 50μg/mL) and clerodin (1.43±0.02 at 40 μg/mL) treated plants (Figure 11& **S**12, **S**12a).

**Figure 11:**
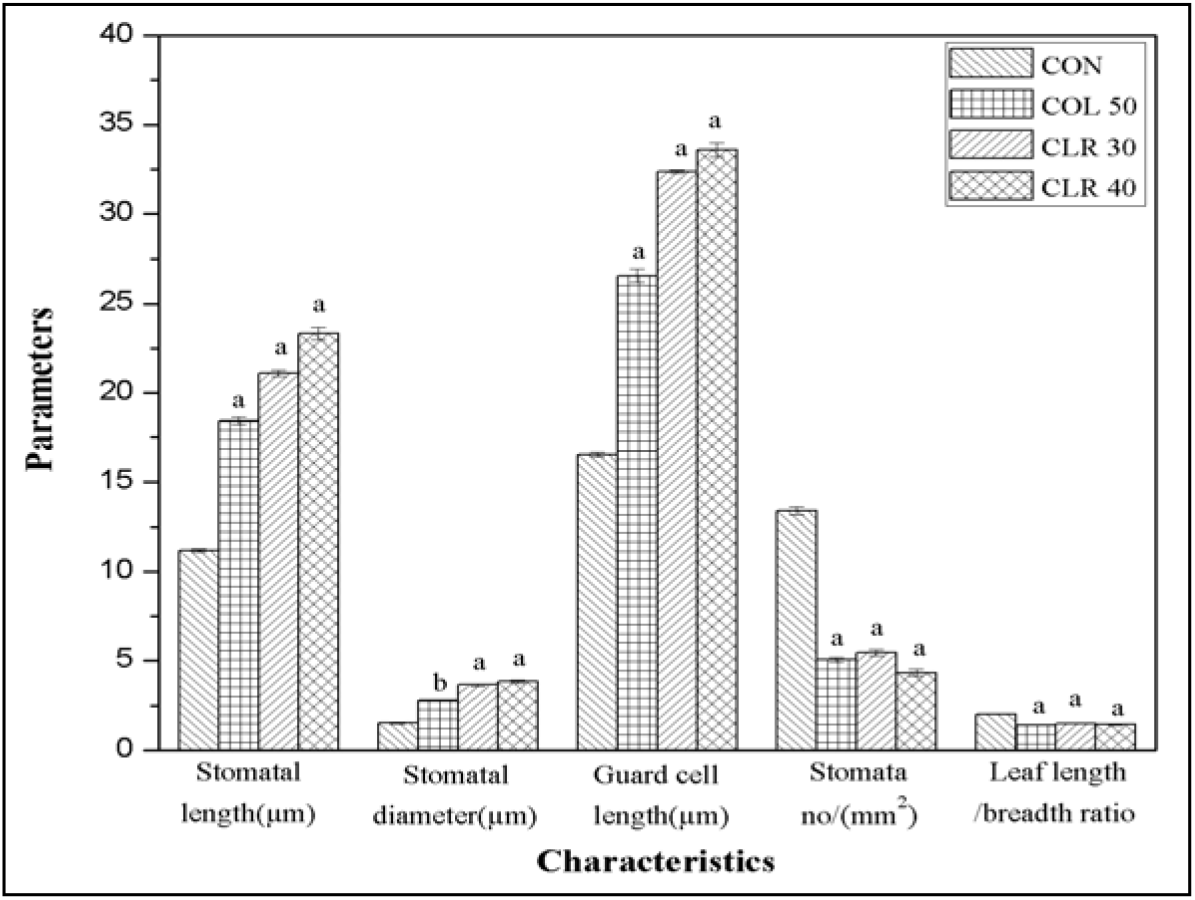
Effect of colchicine and clerodin on morphological characteristics of *Raphanus sativus* L. leaf. Data represented as Mean±SEM. ^a^ significant at p<0.001 and ^b^ significant at p<0.05 as compared to their respective control by Student’s t-test. CON; Control, COL; Colchicine, CLR; Clerodin.

#### 3.5.2. Flow cytometric protoplast DNA content analysis

DNA content analysis through flow cytometry is a convenient method for ploidy determination. The isolated protoplast of colchicine (50 μg/mL) treated plant leaves showed increased polyploidy (41.6 % 2C, 44.4 % 4C, 10.3 % 8C and 2.79 % 16C) and similarly clerodin (40 μg/mL) also induced comparatively more polyploidy (36.4 % 2C, 42.2 % 4C, 17.1 % 8C and 3.23 % 16C) containing cells. Untreated samples show 64.8, 33.2, 1.56, and 0.15 % cells containing respectively 2, 4, 8, and 16 C (Figure 12 & Figure **S12**, A-D).

**Figure 12:**
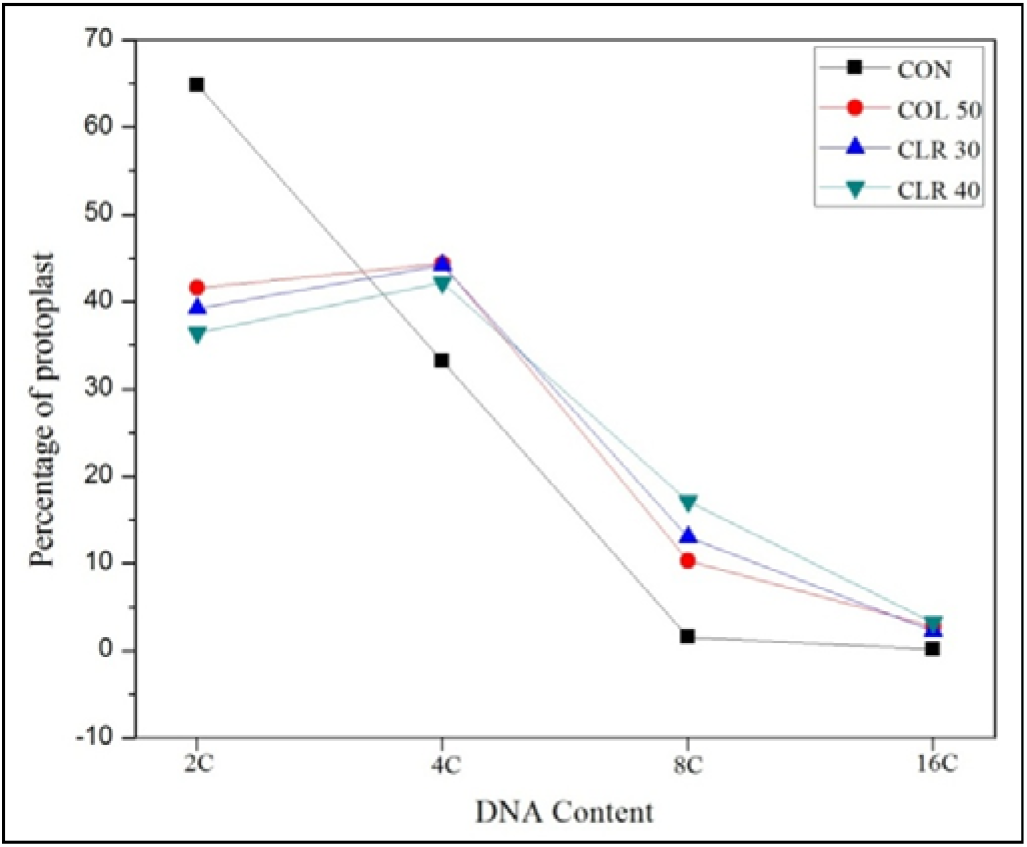
Flow cytometry-based analysis of clerodin and colchicine induced increase in DNA content (polyploidy) in Radish (*Raphanus sativus* L.) protoplast.COL; Colchicine, CLR; Clerodin.

**Figure S12.**
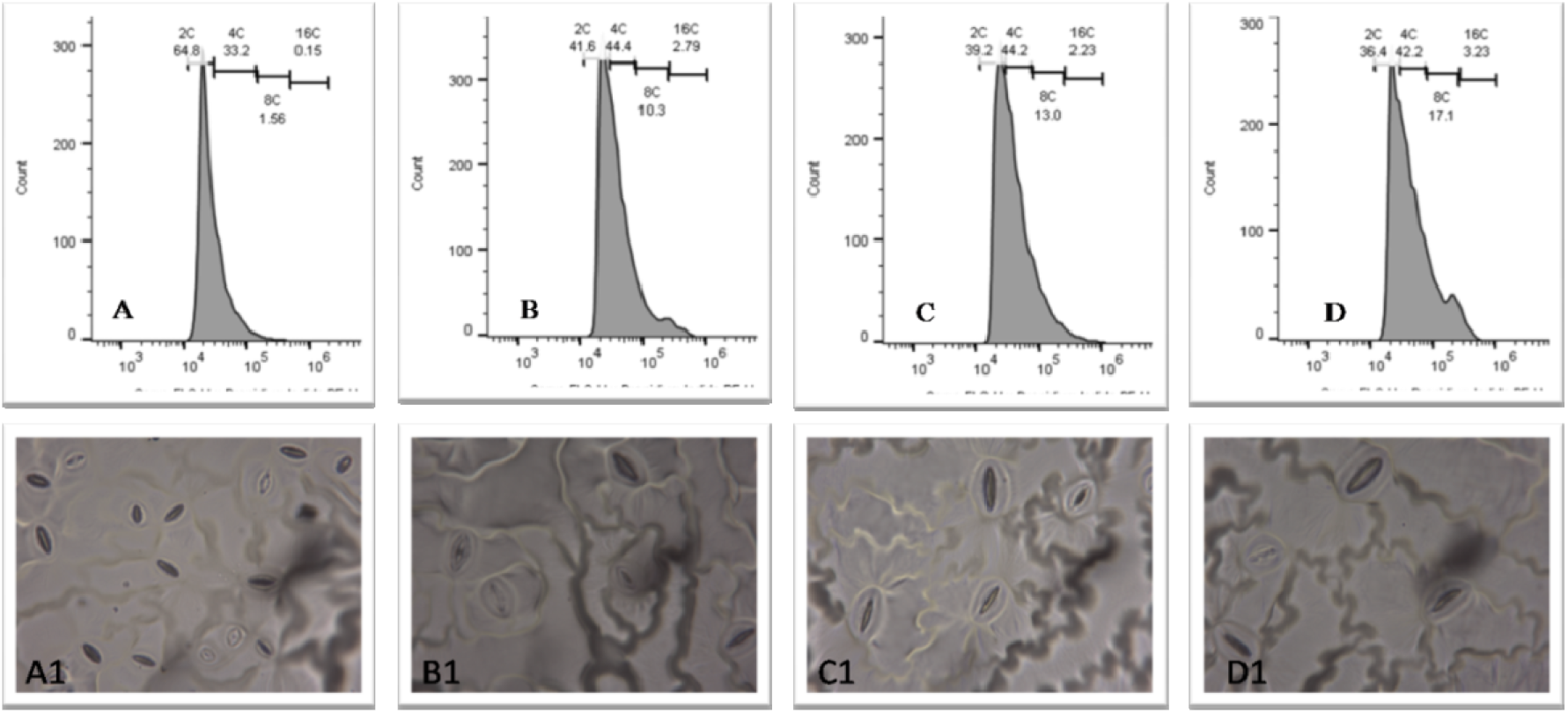
Influence of clerodin and colchicine on flow cytometric assessment of DNA content (A-D), stomatal number and size in radish leaf (A1-D1). A; Untreated, B; colchicine treated (50μg/mL), C; clerodin treated (30 μg/mL), D; clerodin treated (40 μg/mL). A1; untreated, B1; colchicine treated (50μg/mL), C1; clerodin treated (30 μg/mL), D1; clerodin treated (40 μg/mL).

**Figure S12a.**
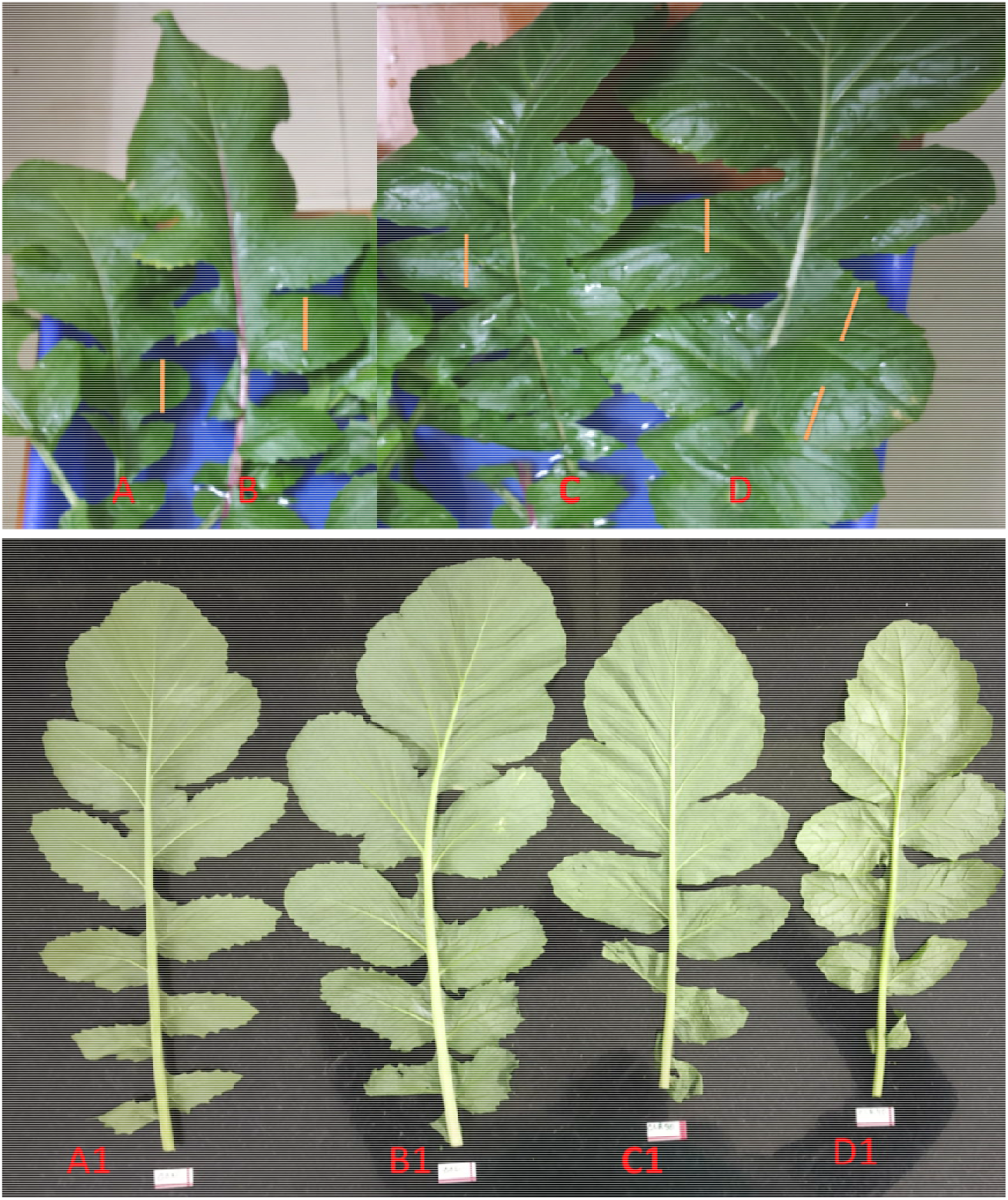
Leaf morphological alteration in colchicine and clerodin induced polyploid Radish. A, A1; untreated leaf, B, B1; colchicine (50 μg/mL) treated leaf. C, C1; clerodin (30 μg/mL) treated leaf, D, D1; clerodin (40 μg/mL). A-D, represent mature leaf and A1-D1 immature leaves of radish.

### 3.6. Molecular docking analysis

Molecular docking results showing the free binding energy of colchicine and clerodin with tubulin along with the interacting residues and estimated inhibition constant, Ki, is tabulated below in Table 3.

**Table 3:**
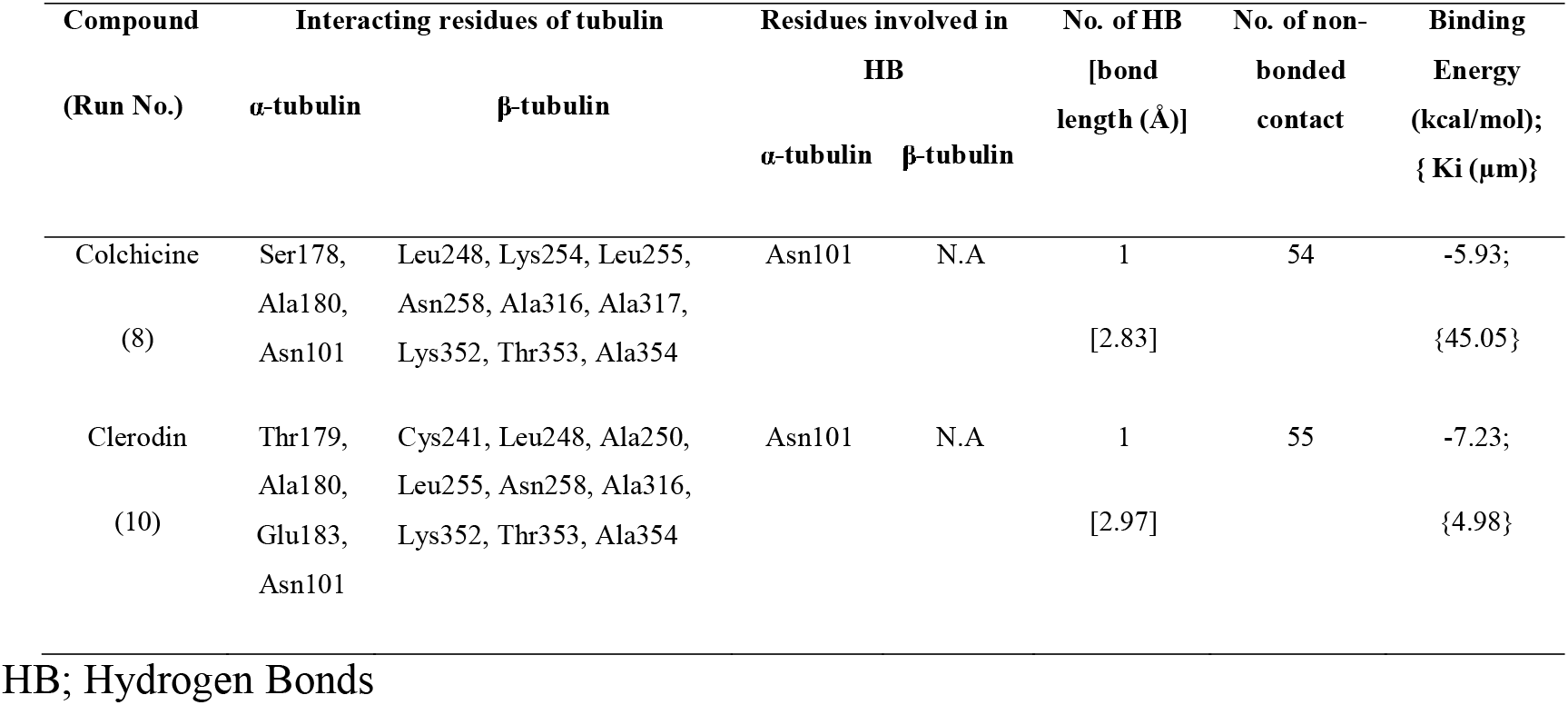
Free binding energy with estimated inhibition constant and interacting residues of Colchicine and Clerodin with tubulin.

It was observed that clerodin represented a better conformation than colchicine and showed to interact with the residues Asn101, Leu248, Leu255, Asn258, Ala316, and Lys352, the amino acid residues that are get interacted with colchicine as well (Figure 13). A more detailed molecular interaction study of colchicine and clerodin with tubulin using LIGPLOT revealed that colchicine interacts with Asn101, Ser178, and Ala180 residues while Clerodin interacted with Asn101, Thr179, Ala180, and Glu183 residues in the α-subunit of tubulin. Both colchicine and clerodin interacted with the residues Asn101 and Ala180 and formed hydrogen bonds of length 2.83Å and 2.97Å respectively with Asn101 and had hydrophobic interaction with Ala180. In the β-subunit of tubulin, the residues Leu248, Lys254, Leu255, Asn258, Ala316, Ala317, Lys352, Thr353, and Ala354 interacted with colchicine and had hydrophobic interaction while clerodin showed hydrophobic interaction with Cys241, Leu248, Ala250, Leu255, Asn258, Ala316, Lys352, Thr353, and Ala354 residues. The residues Leu248, Leu255, Asn258, Ala316, Lys352, Thr353, and Ala354 were common residues forming hydrophobic contacts with both colchicine and clerodin. Figure 13 represents the binding pockets, interacting residues, and LIGPLOT analysis of tubulin for colchicine (A, C, E) and clerodin (B, D, F). The detailed molecular interaction analysis of colchicine and clerodin with tubulin is tabulated in Table 3. Finally, the most favorable conformations of colchicine and clerodin with tubulin were visualized in UCSF Chimera for a better understanding of the binding pockets of colchicine and clerodin (Figure 13). It is clear that clerodin not only binds to the same pocket where colchicine binds with tubulin but also showed a much better conformation having a binding energy of −7.23 kcal/mol compared to that of colchicine having a binding energy of −5.93 kcal/mol.

**Figure 13:**
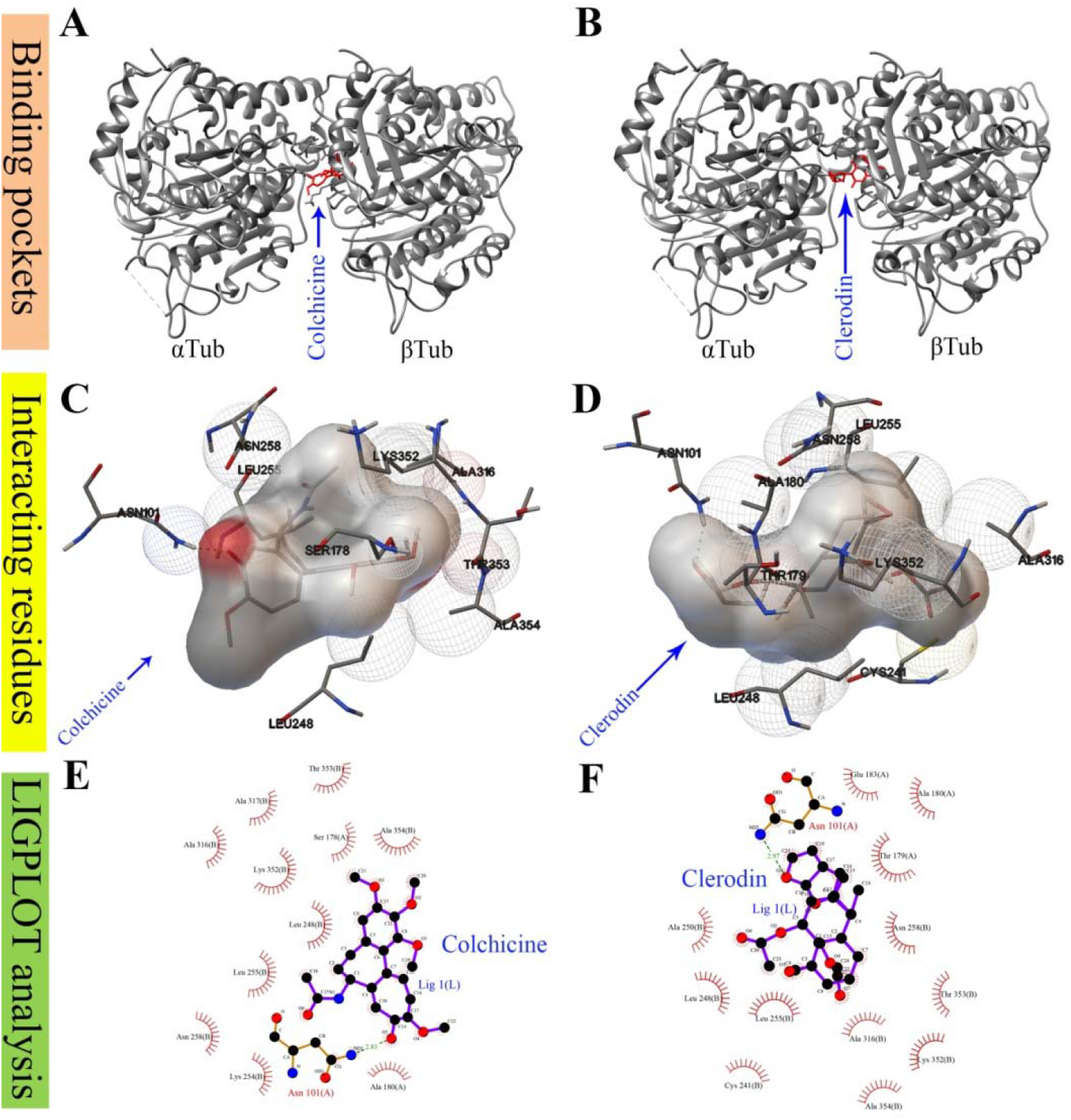
The binding pockets, interacting residues and LIGPLOT analysis of tubulin for colchicine (A, C, E) and clerodin (B, D, F).

## 4. Discussion

The present investigation emphasizes the comparative study between clerodin and colchicine for their microtubule destabilization based pro-metaphase/metaphase arresting, mitotic abnormalities, and polyploidy inducing effects. Earlier studies demonstrated that clerodin is one of the bioactive principles of *C. viscosum* (Barton et al. 1961, Abbaszadeh et al. 2012).

Terpenoids, the largest class of natural products, are well known for their use in the fragrance and flavor industries, in the pharmaceutical and chemical industries (Gershenzon et al. 2007). The present result shows an increase in the frequency of MI in both HPBLs and HEK 293 after clerodin treatment than untreated samples indicating its metaphase arresting and cell cycle delay inducing effects. The observed chromatid condensation and irregular arrangements of chromosomes in prometaphase plate indicate colchicines-like microtubule destabilization effect of clerodin. The present results indicate that HEK 293 cells are more susceptible to clerodin than HPBLs. In this study, the standard drug colcemid was used as a positive control since it has been used to arrest the cell at metaphase in HPBLs for chromosomal aberration analysis (Holden HE, Crider PA, Wahrenburg MG. 1980, Ray and Chatterjee, 2006).

Flowcytometric data show that clerodin has the potential to increase G2-M frequencies in the MCF-7 cell line. Clerodin induces 11.28 and 16.25% of G2/M-phase arrest after treatment with 100 and 200μg/mL, respectively whereas in the untreated cells the frequency of G2/M was 6.10%. The quantitative reduction of DNA content was noticed in G2/M-phase arrest cells which could be due to the loss of chromosomes as micronuclei. Moreover, confocal microscopic tubulin imaging in MCF-7 cells indicates that clerodin induces depolymerization of tubulin network which could bean underlined mechanism for G2/M-phase arrest. Such G2-M-phase arrest by the synthetic compound CC-5079, a potent tubulin polymerization inhibitor, exerts antitumor activity (Zhang et al. 2006).

In this study, clerodin induced root growth retardation, metaphase arrest, mitotic abnormalities, and polyploidy in *A. cepa* root tip cells. With respect to root growth inhibition, clerodin (IC_50_ – 18.98 ± 4.16 μg/mL) has shown more effective than colchicine (IC_50_ – 29.83 ± 2.12 μg/mL).Our data indicate that meristems are sensitive to clerodin and *Allium cepa* root growth retardation may be due to induced mitotic abnormalities and cell cycle arrest as it was described earlier (Fiskesjö 1985; Ray et al. 2012).

Both clerodin and colchicine treatment increased the frequency of mitotic index in a dose-dependent manner in *A. cepa* root tip cells which were reduced after 16 and 32 h of recovery. Reduction in the frequency of prophase, anaphase, and telophase by clerodin and colchicine were observed in all the treated samples. Reduction of mitotic index in recovery samples may occur due to the reconstruction of arrested metaphase nuclei into interphase nuclei (Macleod and O’riordan 1966).

Assessment of mitotic abnormalities indicates that the total aberrant cell frequencies increased with the increasing concentrations of clerodin and colchicine. Their effects on c-metaphase formation are more or less similar, whereas, clerodin induces more c-metaphase at 16 h recovery as compared to colchicine and this observation also indicates the slow release of its effect as the polarity of clerodin is much lower than colchicine. Formation of c-metaphase or colchicines-like metaphase is directly correlated with the microtubule disruption within the cell and the present investigation indicates that clerodin might have colchicine like microtubule destabilizing activity (Bonciu et al. 2018, Özmen 2010). The notion of microtubule destabilization effect of clerodin further supported by the observed results that revealed a dose-dependent increase in vagrant and laggard chromosome, and micronuclei frequencies in *A. cepa* root tip cells. It is an established fact that the consequences of microtubule destabilization based formation of c-metaphases, vagrant chromosomes, laggard chromosomes, and chromatid break lead to the formation of cells with polyploid and micronuclei (Fernandes et al. 2007). It is reported that the disruption of mitotic spindle might inhibit cytokinesis and such inhibition of cytokinesis and reconstruction of nuclei leads to the formation of polyploid cells (Carvalho et al. 2019, Fenech and Crott 2002, Meng. 2003). Spindle disruption may lead to the unequal separation of chromosomes between the daughter nuclei, which in turn lead to the formation of vagrant and laggard chromosomes and contributing to the formation of unequal-sized nuclei (El-Ghamery et al. 2003) or micronuclei. Thus, clerodin induced increased frequency of laggard chromosomes and micronuclei could be due to inhibition of tubulin polymerization which could be the reason for delayed movement of chromosome towards poles.

Treatment with either clerodin or colchicine shows root swelling in *A. Cepa* which could be due to the generation of polyploid/multinucleate cells. Similar swelling patterns demonstrated earlier in onion and wheat root after treatment with colchicine and leaf aqueous extracts of *Clerodendrum viscosum* (Ray et al. 2013). In the present study, both clerodin and colchicine treatment increased the length and diameter of the stomata, decreased the number of stomata per square millimeter of leaf surface, reduced the length and breadth ratio of a leaf, and increased the protoplast DNA content in the leaf of a treated radish plant as compared to untreated controls. These are the indication of polyploidy induction by the treatment since the decrease in length and breadth ratio makes the leaf shorter and wider in shape, which is a characteristic feature of a polyploid plant (Omidbaigia et al. 2010). Moreover, the elevation of DNA content in leaf protoplast and altered leaf morphology after clerodin-treatment indicate its higher potentiality in polyploidy induction than colchicine. This notion is further strengthened through LIGPLOT analysis which confirms that both clerodin and colchicine interact with ASN101 residue by hydrogen bonding. Estimated inhibition constant and free binding energy of clerodin and colchicine with tubulin indicates that clerodin has better free binding energy and inhibition constant compare to colchicine. The present results demonstrate similarity with the earlier report where interaction between tubulin domain and colchicine was reported (Massarotti et al. 2012).

## 5. Conclusion

Clerodin is one of the major bioactive compounds of leaf extract of *Clerodendrum viscosum*. Colchicine like actions of clerodin in terms of microtubule destabilization based mitotic abnormalities and plant polyploidy inducing potentials are explored here for the first time. The experimental data revealed that clerodin has metaphase arresting, microtubule destabilization, and polyploidy inducing ability, somewhat better or comparable with colchicine action. Molecular docking analysis revealed for the first time that both of them interact at the common site of tubulin residue indicating a common mechanism of action. The results also indicate similar cytotoxic potentialities of both clerodin and colchicine even though they belong to different chemical groups. The existing plant polyploidy inducing agent, colchicine alkaloid, may be replaced with a clerodane type diterpenoid, clerodin, for polyploidy induction in plant breeding program and opens up further research to validate its therapeutic applications as an alternative drug.

## Acknowledgments

The authors acknowledge Prof. A. Mukherjee for plant species authentication and the financial support of UGC MRP [F.No.42-563/2013 (SR) dt. 22.3.13] and UGC-SRF [FC(Sc)/RS/UGC/ZOO/2018-19/129, w.e.f. 07.04.2018, dated: 04.02.2019], and the DST-PURSE, DST-FIST, and UGC-DRS-sponsored facilities in the Department of Zoology.

