## Supplemental Tables for "Deciphering colchicine like actions of clerodin in terms of microtubule destabilization based mitotic abnormalities, G2/M-phase arrest, and plant polyploidy"

**SUPPLEMENTARY DATA**

**Table S1:** Effects of clerodin and colcemid on the metaphase percentage in HPBLs and HEK 293 cells.

|  | **H** | **Comp** | **Conc(μg/mL)** | **TC** | **TMC** | **MI%**  **Mean ±SE** |
| --- | --- | --- | --- | --- | --- | --- |
| HPBLs | **3** | Control | 0 | 3792 | 10 | 0.26±0.03 |
|  |  | Colcemid | 0.1 | 3575 | 125 | 3.51±0.18^a^ |
|  |  | Clerodin | 100 | 3695 | 66 | 1.76±0.18^a^ |
|  | **6** | Control | 0 | 4026 | 16 | 0.40±0.02 |
|  |  | Colcemid | 0.1 | 3893 | 179 | 4.59±0.16^a^ |
|  |  | Clerodin | 100 | 4261 | 122 | 2.88±0.19^a^ |
| HEK 293 | 3 | Control | 0 | 3181 | 4 | 0.12±0.03 |
|  |  | Colcemid | 0.1 | 3662 | 174 | 4.74±0.2^a^ |
|  |  | Clerodin | 100 | 3407 | 210 | 6.18±0.33^a^ |
|  | 6 | Control | 0 | 3354 | 6 | 0.18±0.01 |
|  |  | Colcemid | 0.1 | 3791 | 226 | 5.92±0.31^a^ |
|  |  | Clerodin | 100 | 3214 | 344 | 10.62±0.61^a^ |

^a^significant at p<0.001 as compared to their respective control by 2x2 Contingency χ2 -test with respective d.f. = 1, H; Hours, Comp; Compound, Conc; Concentration, TC; Total number of cells, TMC; Total number of metaphase, MI; Mitotic index.

**Table S2:** Root growth retardation effect of clerodin and colchicine on *A. cepa*.

| **Extract fractions**  **(µg/mL)** | | **Root length (cm)** | | | **Growth rate (cm/24H)** | | **Root length retardation percentage (%)** | | | **Growth rate retardation percentage (%)** | |
| --- | --- | --- | --- | --- | --- | --- | --- | --- | --- | --- | --- |
|  |  | **24H** | **48H** | **72H** | **48H** | **72H** | **24H** | **48H** | **72H** | **48H** | **72H** |
| **Control** | | 3.72±0.18 | 5.06±0.16 | 6.28±0.05 | 1.34±0.02 | 1.12±0.13 | 00 | 00 | 00 | 00 | 00 |
| **Clerodin** | **5** | 3.18±0.14^b^ | 4.05±0.11^a^ | 4.86±0.02^a^ | 0.86±0.03^a^ | 0.81±0.13^a^ | 14.37±0.62^b^ | 19.97±0.51^a^ | 22.49±0.82^a^ | 35.61±1.99^a^ | 32.27±2.60^a^ |
|  | **10** | 3.00±0.16^a^ | 3.78±0.10^a^ | 4.25±0.04^a^ | 0.78±0.05^a^ | 0.46±0.06^a^ | 13.28±5.64^a^ | 25.18±0.68^a^ | 32.32±0.66^a^ | 41.64±3.49^a^ | 62.08±0.74^a^ |
|  | **20** | 2.81±0.17^a^ | 3.35±0.09^a^ | 3.63±0.05^a^ | 0.54±0.07^a^ | 0.28±0.04^a^ | 24.49±1.33^a^ | 33.74±0.35^a^ | 42.08±0.85^a^ | 59.83±5.43^a^ | 76.80±2.11^a^ |
|  | **40** | 2.46±0.13^a^ | 2.98±0.10^a^ | 3.19±0.08^a^ | 0.51±0.03^a^ | 0.21±0.03^a^ | 33.77±0.86^a^ | 41.14±0.64^a^ | 49.01±1.14^a^ | 61.74±1.93^a^ | 82.26±1.93^a^ |
|  | **80** | 1.87±0.27^a^ | 2.5±0.11^a^ | 2.62±0.08^a^ | 0.42±0.04^a^ | 0.12±0.03^a^ | 44.05±0.83^a^ | 50.74±0.80^a^ | 58.28±1.24^a^ | 68.30±3.53^a^ | 90.42±1.83^a^ |
| **Colchicine** | **5** | 3.35±0.14^c^ | 4.49±0.11^c^ | 5.52±0.03^a^ | 1.11±0.00^c^ | 1.02±0.12^b^ | 9.88±0.68^c^ | 11.14±0.60^c^ | 12.07±0.47^b^ | 14.70±1.27^c^ | 15.66±0.64^b^ |
|  | **10** | 3.28±0.16^b^ | 4.27±0.12^b^ | 5.09±0.05^a^ | 0.98±0.03^b^ | 0.82±0.14^a^ | 12.39±0.11^b^ | 15.64±0.35^b^ | 18.94±0.72^a^ | 27.34±2.11^b^ | 33.66±4.67^a^ |
|  | **20** | 3.05±0.14^a^ | 3.71±0.12^a^ | 4.07±0.08^a^ | 0.65±0.03^a^ | 0.36±0.11^a^ | 17.97±0.92^a^ | 26.71±1.11^a^ | 35.09±1.02^a^ | 51.01±2.15^a^ | 71.31±6.34^a^ |
|  | **40** | 2.72±0.13^a^ | 3.30±0.09^a^ | 3.56±0.02^a^ | 0.58±0.06^a^ | 0.25±0.07^a^ | 26.75±1.33^a^ | 34.73±0.30^a^ | 43.30±0.00^a^ | 56.80±4.01^a^ | 79.87±4.22^a^ |
|  | **80** | 2.40±0.12^a^ | 2.92±0.13^a^ | 3.12±0.10^a^ | 0.52±0.08^a^ | 0.19±0.03^a^ | 35.53±1.22^a^ | 42.31±0.95^a^ | 50.32±1.52^a^ | 60.94±6.33^a^ | 84.08±1.41^a^ |

Data represented as mean ± sem. Data represented as Mean ± SEM.^a^ significant at p<0.001 and ^b^ significant at p<0.01, ^c^ significant at p<0.05 as compared to their respective control by t-test with respective d.f. = 1.

**Table S3:** Effect of colchicine and clerodin on morphological characteristics of *Raphanus sativus* L. leaf.

| **Characteristic** | **CON** | **COL 50** | **CLR30** | **CLR40** |
| --- | --- | --- | --- | --- |
| **Stomatal length (µm)** | 11.18±0.10 | 18.42±0.23^a^ | 21.10±0.17^a^ | 23.33±0.36^a^ |
| **Stomatal diameter (µm)** | 1.53±0.03 | 2.78±0.05^b^ | 3.65±0.07^a^ | 3.83±0.08^a^ |
| **Guard cell length(µm)** | 16.56±0.14 | 26.56±0.37^a^ | 32.37±0.09^a^ | 33.60±0.39^a^ |
| **Stomatal number/(mm^2^)** | 13.41±0.22 | 5.07±0.14^a^ | 5.46±0.18^a^ | 4.33±0.19^a^ |
| **Leaf length/breadth ration** | 2.02±0.02 | 1.40±0.02^a^ | 1.48±0.01^a^ | 1.43±0.02^a^ |

Data represented as Mean±SEM. ^a^ significant at p<0.001 as compared to their respective control by Student’s t -test. CON; Control, COL; Colchicine, CLR; Clerodin.

3
